# CACNB4 overexpression decreases dendritic spine density in sex-specific manner

**DOI:** 10.1101/2022.02.02.478824

**Authors:** Emily M. Parker, Nathan L. Kindja, Rebecca A. DeGiosio, Ryan B. Salisbury, Josh M. Krivinko, Claire E. J. Cheetham, Matthew L. MacDonald, Robert A. Sweet

## Abstract

The canonical voltage-gated calcium channel (VGCC) subunit complex is comprised of the α1 subunit, the ion permeable channel, plus three auxiliary subunits: β, α2δ and γ. β is the most extensively studied auxiliary subunit and is necessary for proper forward trafficking of the α1 subunit to the plasma membrane. α1 subunits mediate voltage-dependent movement of calcium ions into the cytoplasm of neurons, including at dendritic sites, where increased intracellular calcium initiates signaling cascades that shape structural and functional plasticity of dendritic spines. Genetic studies strongly implicate calcium signaling dysfunction in the etiology of neurodevelopmental disorders including schizophrenia. Dendritic spine density (DSD) is significantly decreased in schizophrenia in primary auditory cortex where DSD is driven by loss of small spines, and small spine loss is associated with increased peptide levels of ALFDFLK found in the VGCC β subunit β4. Overexpessing *CACNB4* to increase β4 levels selectively reduced small spine density in cortical neuron cultures. The studies described herein set out to validate this *in vitro* observation in an intact mammalian system within a neurodevelopmental context. We overexpressed *CACNB4* in neurodevelopment and assessed DSD and morphology in cerebral cortex of male and female mice at an adult timpoint. We then characterized β protein levels and β4 protein-protein interactions in male and female mouse cortex. Overexpression selectively reduced small dendritic spine density but this effect was present only in female mice and did not appear to result from estrous stage. Instead, the sex-dependent effect on DSD corresponded to sex differences in the β4 interactome of male versus female mice: the VGCC β subunit β1b was significantly enriched in the β4 interactome of brain tissue of male mice, and thus may have served to mitigate VGCC overexpression-mediated spine loss in male mice.

## INTRODUCTION

β is the most extensively studied auxiliary subunit of voltage-gated calcium channels (VGCCs)^1^. β, along with two other auxiliary VGCC subunits, α2δ and γ, typically binds to the α1 (ion permeable channel) VGCC subunit with 1:1:1:1 reversible stoichiometry^2–4^. There are four β protein subfamilies, β1-4^5^. Each β protein subtype is encoded by a separate gene with multiple splice variants, all of which are highly expressed at transcript and protein levels in mouse brain except β1a, β1c-d, β2d, and β2e^2,6^. β4a expression appears limited to the cerebellum^2^. β subunits predominate in neuronal subcellular compartments depending on the high-voltage activated (HVA) VGCC α1 subunits they preferentially bind to, traffick and modulate^2^. Although the binding affinity of β to α1 is high, ranging from K_D_ of 2–54nM, α1 to β pairing is reversible and α1-β reshuffling may occur either to compensate for loss of β or through competitive replacement^3,4,7–9^.

β4 is the focus of the current study and is encoded by the CACNB4 gene, which has now five known splice variants^10^. β4 variants are generally expressed highly in brain, including in rodents, where transcript and protein levels are altered in several brain regions as a function of age^2,11,12^. As a general rule, β4 preferentially binds the presynaptic α1 Ca_V_2.1 (P/Q-type) VGCC^13,14^. β4 has also been shown to bind to approximately 40% of α1 Ca_V_1 (L-type) VGCCs, in addition to other calcium channels: Ca_V_2.2 (N-type) and Ca_V_2.3 (R-types)^2,15,16^. Neuronal subcellular distribution of β4 appears diffuse; β4 is located at the plasma membrane, in intracellular space, in dendrites and dendritic spines and in axons. Electron microscopy revealed murine cerebellar and hippocampal β4 levels are significantly higher in intracellular space than other compartments^12^. β4 co-localizes with VGLUT1 in presynaptic terminals of glutamatergic neurons and is present in synapse preparations for mass spectrometry^10,17^. β4b is the only isoform specifically demonstrated to exibit nuclear targeting^18^.

β proteins (β1-4) perform similar, non-identical functions^2^. β is considered the most critical auxiliary VGCC subunit and work over a number of years has demonstrated β’s most important function: forward trafficking the α1 subunit of HVA VGCCs toward the plasma membrane^19–22^. Once established in the plasma membrane, α1 subunits mediate voltagedependent movement of calcium ions down a steep gradient, from extracellular sites into the cytoplasm of neurons and cardiomyocytes, among others^23–25^. Depolarizations generated at synapses and back-propagating action potentials trigger VGCC-mediated calcium influx in dendrites^23^. Calcium, upon entering neurons, initiates signaling cascades, a number of which shape structural and functional plasticity of dendritic _spines_^23,26,27^.

Genetic studies strongly implicate calcium signaling dysfunction, including alterations to VGCC subunit signaling in particular, in the etiology of epilepsy and neurodevelopmental disorders, among these schizophrenia^28,29^. Dendritic spine density is significantly reduced in primary auditory cortex (A1) in schizophrenia and loss of small spines drives this reduction^30–34^. Follow-up analysis of postmortem A1 brain tissue from individuals with schizophrenia and non-psychiatric controls revealed levels of the tryptic peptide ALFDFLK found in the β4 calcium channel subunit inversely correlated with dendritic spine density of small, but not large, spines. Further, CACNB4 overexpression (abbreviated here: β4OE) selectively reduced small spine density in dissociated primary cortical neuron cultures fixed on day *in vitro* 15^34^.

The goal of the current study was to validate the *in vitro* observation in an intact mammalian system. To accomplish this we assessed dendritic spine density (DSD) and morphology in the cerebral cortex of male and female mice at the adult timepoint of postnatal day 84 (P84) following CACNB4 overexpression (β4OE). We further characterized β protein levels and β4 protein-protein interactions in murine cortex. We found that β4OE selectively reduced density of small dendritic spines, but the effect of β4OE on spines was present only in female mice. This female-specific effect did not appear to result from estrous stage. Instead, the sex-dependent effect of β4OE on DSD corresponded to sex differences in the β4 interactome. Of note, the β1b VGCC subunit was significantly enriched in the β4 interactome of brain tissue from male, relative to female C57BL/6J mice, and thus may have served to mitigate spine loss in β4OE male mice. β1b is the β1 isoform expressed in brain.

## MATERIALS & METHODS

### Experimental Animals

#### *AAV-exposed* C57BL/6J *mice*

All experiments were approved by the IACUC at the University of Pittsburgh in accordance with the guidelines outlined in the USPHS Guide for Care and Use of Laboratory Animals. E16 pregnant C57BL/6J dams were acquired from The Jackson Laboratory (Bar Harbor, ME) and singly housed in BSL-2 biocontainment in standard microisolator cages (Allentown Caging Equipment, Allentown, NJ) on a 12h light/dark cycle with food and water provided ad libitum. The Synapsin-driven mCherry fluorescent adeno-associated virus (AAV) AAV2-hSyn-mCherry (Titer≥5×10^12^vg/ml), which confers long-term transgene expression of mCherry in neurons, was obtained from AddGene (#114472-AAV2). Two other AAVs were acquired, both from Penn Vector Core: AAV2-CaMKII-eGFP-WPRE (Titer=1.088e13gc/ml) and AAV2-CaMKII-CACNB4-P2A-eGFP-WPRE (Titer=4.94e13gc/ml). Two versions of AAV injectate were prepared by adding 1μL AAV2-hSyn-mCherry to either 1μL control AAV (AAV2-CaMKII-eGFP-WPRE) or 1μL AAV-CACNB4 (AAV2-CaMKII-CACNB4-P2A-eGFP-WPRE) diluted in sterile filtered 1x phosphate buffered saline (PBS) at 1:10^35,36^. Diluted AAV was used to achieve sparse AAV transduction. Control (CN) mice were produced by exposure to injectate containing 1μL control AAV. β4 overexpression (β4OE) mice were produced by exposure to 1μL AAV-CACNB4 injectate. AAV2-hSyn-mCherry was used to avoid biasing DSD measurement due to differences in GFP fluorescence produced by the control versus AAV-CACNB4.

Dendritic spine study design and execution schema is found in **Supplemental Figure 1**. AAV solutions were randomly coded “A” and “B” prior to injection, blinding investigator to group. ~50% of each litter was exposed to injectate A and 50% to B. In total, twenty four P0-P2 C57BL/6J mouse pups were exposed to injectate A or B using the bulk regional AAV injection (BReVI) procedure^37^. Neonates were cryoanesthesized^38^ to induce brief hypothermia until response to toe pinch was absent. AAV solution was injected intracranially 1mm rostral to the left earbud and 1mm lateral from the midline using a 1mL Luer-lock syringe connected to a pulled glass micropipette with a sharp tip. Toe amputation was performed while pups were anesthetized for group tagging. Mouse pups were returned to home cage following thrombus at site of toe amputation, waking and 10-12min rewarming. Mice were housed with littermates and dam following the BReVI procedure until 3-weeks following birth (P21), at which point they were weaned and housed with same-sex littermates until P84. Each cage of weaned animals was provided environmental enrichment (hut and exercise wheel) starting at P21, in accordance with Institutional Animal Care and Use Committee (IACUC) policies at the University of Pittsburgh. AAV-mediated β4 overexpression using the constructs described above was verified using western blot of three female C57BL/6J mice (1 AAV-naïve, 1 β4OE and 1 CN) (**Figure 1A**); SDS-PAGE and western blot methods are described below. Experiments were approved by the IACUC at the University of Pittsburgh in accordance with the guidelines outlines in the USPHS Guide for Care and Use of Laboratory Animals.

**Figure 1.**
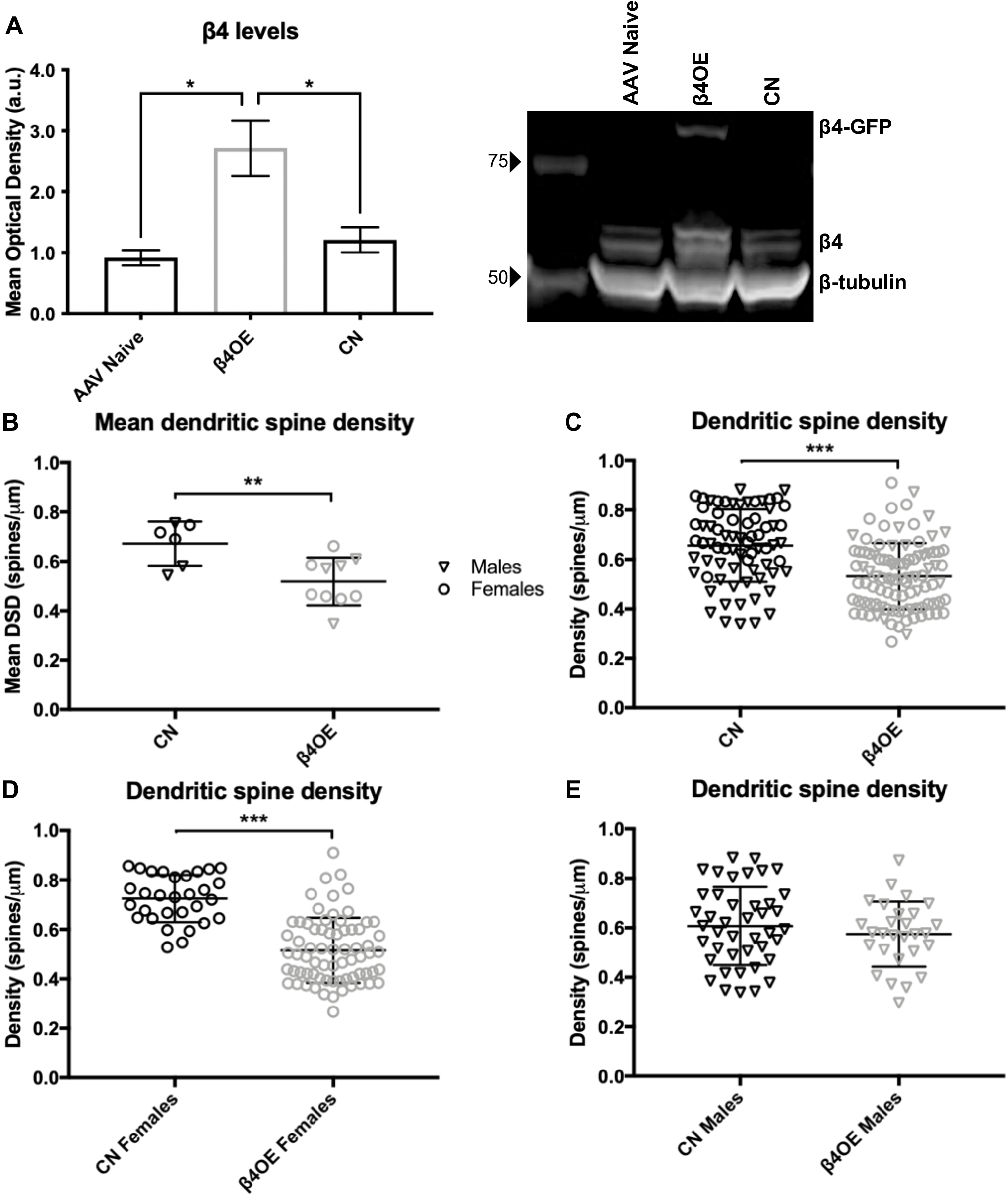
β4 overexpression (β4OE) significantly reduced dendritic spine density (DSD) on minor basal branches of layer 5 (L5) pyramidal cells in sensory cortex. All figures: n.s. p>0.05, * p<=0.05, ** p<=0.01, *** p<=0.001. **A)** Western blot confirming β4 overexpression. (Left) Mean optical density (a.u.) of β4 was significantly increased in β4OE compared to AAV Naïve (t=3.818, DF=2, p=0.0311) as well as to CN (t=3.016, DF=2, p=0.0473). Error bars = SEM. (Right) Western blot showing β4 bands at ~51-55kD as well as band at ~82kD in β4OE tissue, which is the predicted molecular weight of a β4-GFP fusion protein (GFP molecular weight = 27kD). Mean optical density of β4-GFP fusion protein band was not included in assessment of overall β4 overexpression. **B)** β4OE significantly reduced mean DSD at mouse level (F=9.249, DF=3, p=0.01). Data points are mean DSD values from individual mice. Error bars = SD. **C)** DSD was significantly reduced in β4OE, compared to control (CN) mice at the neuron level (F=30.922, DF=3, p<0.001) (n=42 from CN males, n=28 from β4OE males, n=30 from CN females and n=70 from β4OE females). Main effect of sex on DSD was not significant (F=1.812, DF=3, p=0.180). There was a significant genotype by sex interaction (F=16.372, DF=3, p<0.001). Data points are DSD from individual neurons. Error bars = SD. **D)** β4OE significantly reduced DSD in neurons from female mice (F=61.899, DF=1, p<0.001) **E)** In contrast, β4OE did not significantly impact DSD in neurons from male mice (F=0.838, DF=1, p=0.363). Data points in **D** and **E** are DSD from individual neurons. Error bars = SD.

#### Estrous Stage Assessment

Stage of estrous cycle on day of sacrifice was determined for AAV-injected female CN and β4OE mice by evaluating vaginal cytology prior to quantitative microscopy. Vaginal lavage was performed on each female mouse just prior to anesthesia. Briefly, 20μL ddH2O was dispensed onto the vaginal canal opening, fluid drawn back up into a pipette tip and transferred to a glass slide. After air drying, the slide was examined at 20x on a light microscope. Estrous stage (proestrus, estrus or metestrus/diestrus) was estimated based on ratio of cornified squamous epithelial cells, leukocytes and/or nucleated epithelial cells in each sample^39,40^.

#### AAV-naïve C57BL/6J mice

Protein assays were performed to characterize β protein levels and the β4 interactome in brain tissue from male and female wildtype mice^41^. For SDS-PAGE and western blot, four male P28, four female P28, four male P84 and four female P84 mice were anesthetized with a lethal intraperitoneal injection of Nembutal (150mg/kg) and transcardially perfused with ice-cold normal saline. Brains were extracted and rapidly frozen. Bilateral cortical tissue from each brain was separated from the cerebellum and cortex anterior to A1. For CoIP-MS, four additional male P84 and four additional female P84 C57BL/6J mice were sacrificed by lethal CO2 followed by rapid decapitation. Brains were extracted, rapidly frozen and stored at −80°C until immunoprecipitation.

### Quantitative Microscopy

#### Perfusion and Tissue Processing

Twenty CN and β4OE mice were euthanized at P84: 4 male CN, 6 male β4OE, 4 female CN, and 6 female β4OE mice. Mice were weighed, deeply anesthetized with a lethal intraperitoneal injection of Nembutal (150mg/kg) and transcardially perfused with ice-cold 1x PBS followed by 4% paraformaldehyde (PFA). Brains were rapidly extracted and postfixed in 4% PFA for 24h and then moved to 18% sucrose for 24h and stored at −30°C in 30% ethylene glycol and 30% glycerol in phosphate buffer (cryoprotectant) until sectioning. 60μm-thick coronal tissue sections were cut on a cryostat into 12-well plates containing cryoprotectant and placed in −30°C for long-term storage.

#### Immunohistochemistry

Free-floating tissue sections from CN and β4OE mice corresponding to plates 55, 57, 59 and 61 in The Mouse Brain In Stereotaxic Coordinates^42^ (−2.92mm, −3.16mm, −3.4mm and −3.88mm from bregma, respectively) were selected for immunohistochemistry. The region of interest (ROI) for spine analysis is referred to here as sensory cortex and comprises the following adjacent cortical regions: primary and secondary auditory (A1, A2) and visual (V1, V2) cortices, and temporal association cortex (TeA) as defined in the mouse atlas and in Parker et al. 2020^42,43^. Four free-floating sections per mouse were washed in 0.1M PBS, then incubated for 30m in 1% NaBH4 to reduce autofluorescence. Sections were blocked for 3h in a solution of 1% normal goat serum, 3% Triton X-100, 1% bovine serum albumin, 0.1% lysine and 0.1% glycine. Then sections were incubated in primary antibodies guinea pig anti-NeuN (Millipore #ABN90 lot:2834791, 1:2000) and rabbit anti-RFP (Rockland #600-401-376 lot: 39670, 1:1000), for 24h and 96h respectively. Following primary antibody incubation, sections were washed and incubated in secondary antibodies goat anti-guinea pig 405 (Abcam #Ab175678 lot:1972783, 1:500) and goat anti-rabbit Alexa Fluor 568 (ThermoFisher #A11036 lot:997761, 1:500). After 24h incubation in secondary antibodies, tissue sections were washed in a final series and mounted on TruBond 380 micro slide glass (Matsunami, Osaka, Japan) using ProLong Gold antifade mountant (Invitrogen, Carlsbad, CA, USA).

#### Sampling and Confocal Imaging

Images of mounted tissue sections from CN and β4OE mice were captured using an Olympus BX51 WI upright microscope (Center Valley, PA) with an Olympus spinning disk confocal, Hamamatsu ORCA R2 CCD camera (Bridgewater, NJ), BioPrecision2 XYZ motorized stage with linear XYZ encoders (Ludl Electronic Products Ltd., Hawthorne, NY), Lumen 220 light source (Prior Scientific, Cambridge, United Kingdom), excitation and emission filter wheels (Ludl Electronic Products Ltd.) and a Sedat Quad 89000 filter set (Chroma Technology Corp., Bellows Falls, VT), controlled by SlideBook 6 software (Intelligent Imaging Innovations). 2-D images of each tissue section were acquired using an Olympus PlanAPO 1.25x/0.04 N.A. objective and epifluorescent 405nm and 568nm excitation. The mouse atlas^42^ and examination of NeuN and mCherry fluorescence in the 2-D images were used to estimate the laminar and regional location (A1, A2, V1, V2 or TeA) of the cell body of each L5 pyramidal cell imaged (**Supplemental Figure 2A**)^44–46^. L5 mCherry+ pyramidal cells with somal GFP fluorescence (GFP(+)mCherry(+) cells) were identified using an Olympus UPlanSApo 10x/0.40 N.A. objective and captured in 2-D at fixed exposure times (405nm=100ms, 488nm=1500ms and 568nm=150ms at 1.5% Neutral Density)(**Supplemental Figure 2B**). GFP(+)mCherry(+) cells (n=170) were each subsequently captured in a 3-D image stack using an Olympus PlanApo N 60x/1.40 N.A. oil immersion super-corrected objective with spinning disk unit engaged. **Supplemental Figure 2C** shows a single 2-D plane of a 60x stack. mCherry+ L5 pyramidal cells (n=86) lacking GFP (referred to here as β4OE-) in tissue sections from β4OE mice were also captured in 3-D image stacks and serve as the within mouse internal control for neurons of female mice in the β4OE group. Each 60x 3-D stack is a capture site comprised of the cell body of one L5 pyramidal cell and all of its corresponding basal dendrites visible within the 1024×1024 pixel capture window (see also Figure 3C in Parker et al 2020^43^). Neutral Density filter and exposure time for the 568nm channel were optimized for one randomly selected minor basal dendritic segment at each site for each 3-D stack captured. Minor basal dendritic segments were defined as any dendritic segment branching directly off a major or primary basal dendrite. Total tissue thickness was estimated by measuring anti-NeuN labeling in the z-dimension. 3-D image stacks were acquired through the entire thickness of the tissue (0.25μm between each z-plane).

**Figure 2.**
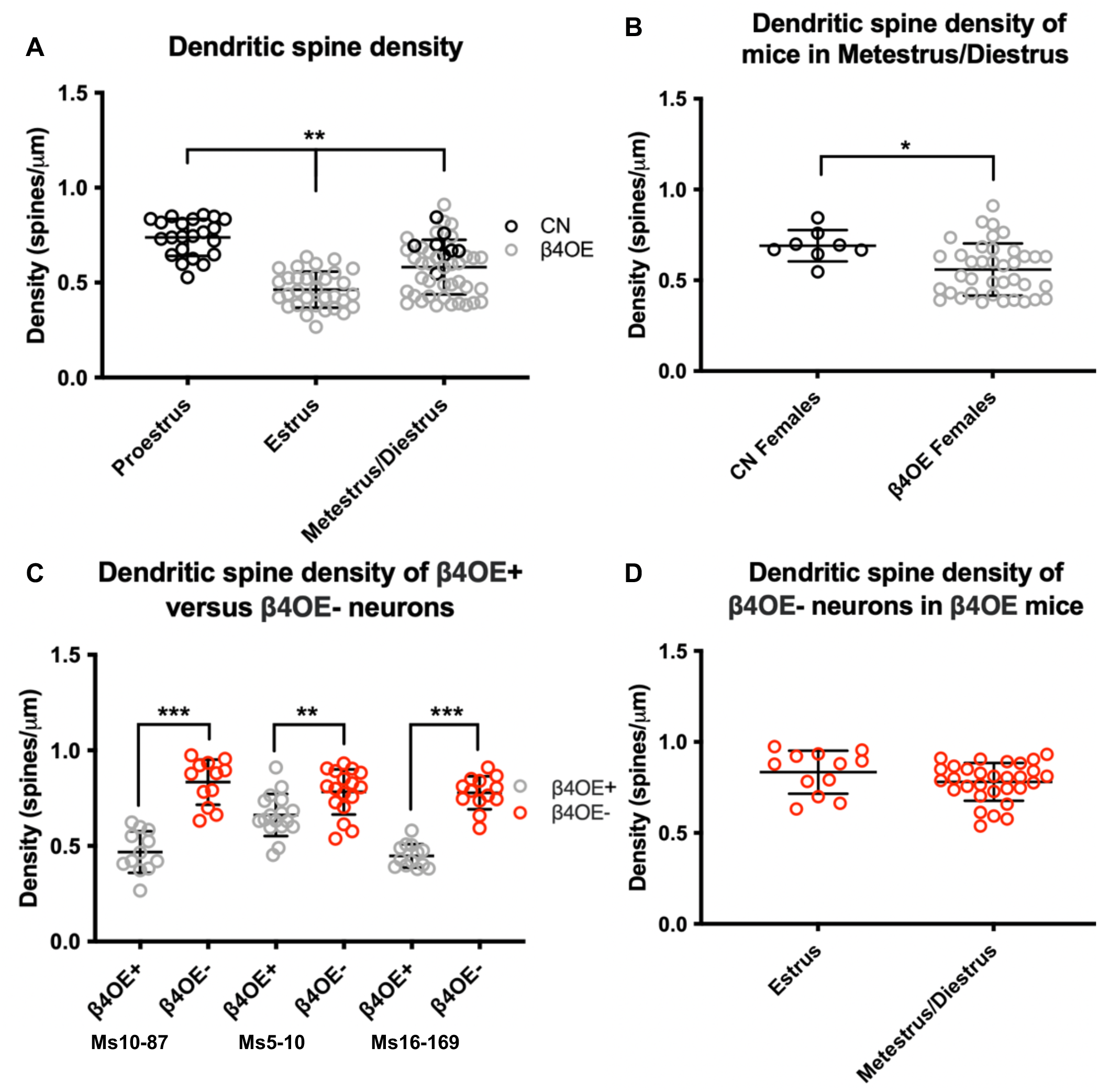
β4OE significantly reduced DSD in female mice regardless of estrous stage. **A)** Estrous stage breakdown of DSD of neurons from female CN and β4OE mice. Spine density was significantly different based on estrous stage (F=6.450, DF=2, p=0.002). Importantly, note both genotypes are represented in metestrus/diestrus, not proestrus, estrus. Data points are from individual neurons. Error bars = SD. **B)** β4OE significantly reduced DSD in female mice in metestrus/diestrus (F=6.190, DF=1, p=0.017). Data points are from individual neurons. Error bars = SD. **C)** Within mouse internal control comparison. DSD of β4OE+ neurons was significantly lower than DSD of β4OE- internal control neurons in three β4OE mice overall (F=103.646, DF=1, p<0.001). There was a significant interaction between estrous stage and condition (β4OE+ versus β4OE-) (F=8.105, DF=1, p=0.006), however main effect of estrous stage was not significant (F=0.985, DF=1, p=0.324). Mice were in different estrous stages (L-R): Ms10-87 was in Estrus (F=67.688, DF=1, p<0.001), and Ms5-10 (F=9.502, DF=1, p=0.004) and Ms16-169 (F=126.518, DF=1, p<0.001) were in metestrus/diestrus on day of sacrifice. Data points from individual neurons. Error bars = SD. **D)** DSD of β4OE-internal control neurons in the three β4OE mice (in **C**) did not significantly differ based on estrous stage (F=2.108, DF=1, p=0.154). Data points from individual neurons. Error bars = SD.

**Figure 3.**
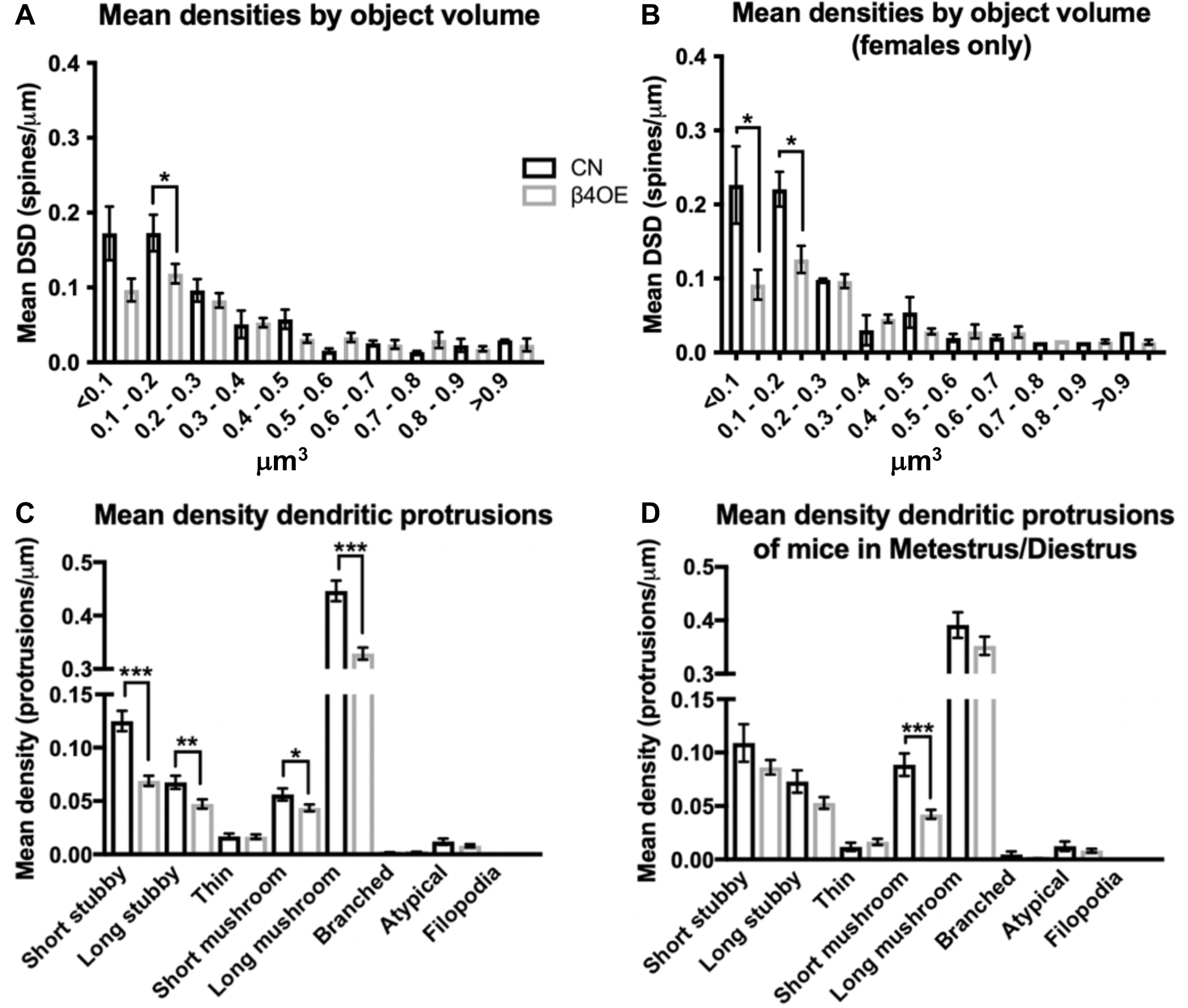
β4OE selectively reduced mean density of spines with small volumes and decreased density dendritic protrusions in female mice. **A)** Mean DSD of small < 0.1μm^3^ objects was significantly reduced in neurons from female β4OE mice, relative to CN (F=5.276, DF=1, p=0.035). Mean DSD of larger objects with volumes > 0.1μm^3^ did not significantly differ based on genotype in female mice. Error bars = SEM. **B)** Focusing exclusively on female mice in metestrus/diestrus, β4OE significantly decreased mean DSD of small objects of 0.1μm^3^ - 0.2μm^3^ volume (F=6.748, DF=1, p=0.032). Error bars = SEM. **C)** β4OE significantly reduced mean density of short stubby (F=33.939, DF=1, p<0.001), long stubby (F=7.043, DF=1, p=0.009), short mushroom (F=4.342, DF=1, p=0.040) and long mushroom (F=30.354, DF=1, p<0.001) spines in female mice (all estrous stages represented) after Bonferroni correction. Collectively, spines in these four categories made up 95.19% of all protrusions observed in female mice. Error bars = SEM. **D)** Focusing exclusively on female mice in metestrus/diestrus, β4OE significantly decreased mean density of short mushroom spines (F=20.160, DF=1, p<0.001). Error bars = SEM.

#### Image Processing and Analysis

SlideBook 6 and Stereo Investigator (MBF Bioscience) software were used for image processing and analysis. 1024×1024 image stacks were first transformed using a no-neighbors smoothing algorithm in SlideBook 6. mCherry fluorescence was used to determine dendrite segment lengths and assess spines in CN and β4OE mice. Minor basal dendritic segments >50μm in length, located >10μm away from the cell body and >3μm from a dendrite branch point were identified in 1024×1024 image stacks and cropped into individual image stacks containing one minor basal dendritic segment each. Mean (SD) distance from the soma of dendritic segments captured for GFP(+)mCherry(+) cells was 15.85 (4.48) μm, indicating these dendritic segments were on proximal branches. Mean distance from soma did not significantly differ by group nor sex (data not shown). Dendritic segment lengths were measured in SlideBook 6 using the line tool. Individual dendritic segments were exported as TIFF series from SlideBook 6 and viewed in Stereo Investigator for spine counting and categorization. Spine objects were manually counted in Stereo Investigator and spine density for each neuron was calculated using the equation: 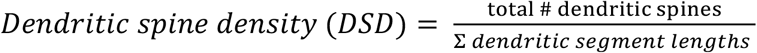

Two neurons per CN or β4OE mouse were randomly selected (from among all neurons that had DSD within 2 standard deviations of the group mean) for further assessment to determine if the significant main effect of group on mean DSD could be driven by a significant difference in density of spines of a particular volume. Spine objects included in the neuron DSD calculation were marked, files coded to blind investigator to neuron and group/sex, and each spine (n=1543 objects) was manually masked using the brush tool in SlideBook 6 (**Supplemental Figure 2D**). Object volume (μm^3^) statistics were extracted for each mask in SlideBook 6 using mask statistics. Spine objects were organized into 10 size bins based on volume as in MacDonald et al 2017^34^. The bin with the smallest spine objects comprised objects of volumes < 0.1μm^3^, the next bin of spines volumes from 0.1-0.2μm^3^ and so on. The bin with the largest spine objects comprised objects measuring > 0.9μm^3^. Dendritic protrusions were classified into one of eight types in Stereo Investigator. Types included short stubby, long stubby, short mushroom, long mushroom, thin, branched or atypical dendritic spine or filopodia. Category was determined based on morphological characteristics described previously (also see **Supplemental Figure 2E**)^43,47,48^

### Protein Assays

#### SDS-PAGE & Western Blot

Two female P0 C57BL/6J mouse pups were exposed to AAV using the BREVI method. One pup was exposed to AAV2-CaMKII-B4-eGFP-WPRE and the second injected with control AAV. On P21 these mice plus one injection naïve female C57BL/6J were all three anesthetized with a lethal intraperitoneal injection of Nembutal (150mg/kg) and transcardially perfused with ice-cold normal saline. Brains were extracted and rapidly frozen. Bilateral cortical tissue from each brain was separated from the cerebellum and cortex anterior to A1. This tissue was then combined with ice-cold Tris-HCl (pH 7.4) and homogenized in lysing tubes in a bead mill. The samples were transferred to new tubes, combined with 2% SDS, probe sonicated, vortexed for 10min at room temperature at 1400RPM followed by centrifugation for 10min at 14,000g. Protein concentration was estimated using a Micro BCA Protein Assay Kit (ThermoFisher Scientific #23235) (R^2^ values > 0.99 for each sample) and protein levels resolved using 1x LI-COR Protein Loading Buffer (Li-Cor #928-40004) loaded into a 4-20% SDS-PAGE gradient gel (ThermoFisher Scientific #26224) and separated for 2h at room temperature in 1x SDS running buffer (Pierce 20x Tris Hepes SDS Buffer #28368) at 75V. Samples were transferred to a polyvinylidene fluoride membrane (Millipore Immobilon-FL PVDF #PFL00010) in 1x Tris Glycine Blotting Buffer (Pierce #28363) for 50m at 4°C at 85V. Non-specific binding was reduced by incubating the membrane in Odyssey Li-Cor Blocking Buffer (Li-Cor #927-4000) diluted in 1x tris buffered saline (TBS) for 1h.The membrane was incubated overnight in Pierce SuperBlock Blocking Buffer (Pierce #37353) with 0.1% Tween 20 (Sigma #P7949) and validated primary antibodies: mouse anti-Ca_V_β4 (Neuromab #75-054, 1:1000) and rabbit anti-β-tubulin (Abcam #ab6046, 1:600,000). The membrane was then incubated in LiCor Blocking Buffer diluted 1:1 with TBS (0.1% Tween 20 and 0.02% SDS) and LiCor IRDye secondary antibodies: goat antimouse 800 (#926-32210, 1:10,000) and goat anti-rabbit 680nm (#926-68071, 1:10,000). β4 isoforms are present at bands ~37.5-59kD. Signal intensity was calculated by normalizing the optical density of β4 to β-tubulin for each technical replicate.

β protein levels in male and female C57BL/6J wildtype mouse A1 were assessed at 2 developmental time points spanning the developmental period of interest for schizophrenia onset^43,49,50^: P28, which is associated with murine estrous cycle commencement (~P24-P30) and P84, corresponding to the age of tissue harvest for AAV-exposed mice. Four male and four female mice in each age group were anesthetized and brains extracted as described above. Mouse A1 tissue was then manually cut from bilateral cryostat sections. A1 tissue was combined with SDS extraction buffer (0.125-M Tris-HCl [ph 7], 2% SDS and 10% glycerol), centrifuged, protein was extracted, and protein concentration estimated using the methods described above. 20μg protein was combined with 1x LI-COR Protein Loading Buffer (10% mercaptoethanol), loaded into an SDS-PAGE gradient gel, separated, transferred to a PVDF membrane and blocked as described above. β-tubulin (50kD) served as loading control. The membrane was incubated overnight in Pierce SuperBlock Blocking Buffer and validated primary antibodies: either mouse anti-Ca_V_β1 (Neuromab #73-052 1:100), rabbit anti-Ca_V_β3 (Alomone Labs #AAC-008, 1:800) or mouse anti-Ca_V_β4 (Neuromab #75-054, 1:1000) and rabbit anti-β-tubulin (Abcam #ab6046, 1:600,000). Membranes were then incubated in LiCor Blocking Buffer as described above and LiCor IRDye secondary antibodies: either goat anti-mouse 800nm (#926-32210, 1:10,000) and goat anti-rabbit 680nm (#926-68071, 1:10,000) or goat anti-rabbit 800nm (#926-32211, 1:10,000) and goat anti-mouse 680nm (#926-68070, 1:10,000). β1 isoforms are present at bands ~37.5-59kD. β3 isoforms are present at bands ~37.5-59kD. β4 isoforms are present at bands ~37.5-59kD. Four biological replicates (4 mice) per group and 1 technical replicate (2 lanes) per mouse were run. Four blots were run for each β protein assessed (β1, β3, and β4), resulting in 12 total blots run. Signal intensity for each mouse was calculated by normalizing the optical density of β1, β3 or β4 to β-tubulin for each technical replicate and then averaging the normalized optical density of the technical replicates. Mean optical density for each group was the average mean β1, β3 or β4/mean β-tubulin of each mouse in the group.

#### Co-immunoprecipitation

Anti-Ca_V_β4 (Neuromab #75-054) was used for β4 immunoprecipitation, having been previously used for western blot (see above) and to immunoprecipitate β4 from synaptosome preparations by a different research group^17^. Additionally, we demonstrated this antibody successfully immunoprecipitated β4 from cortical homogenate in a coimmunoprecipitation (CoIP) western blot assay using goat anti-CACNB4 (Everest Biotech #EB06591, 1:1000) and a donkey anti-goat 800 secondary antibody (LiCor #926-32214, 1:10,000) for detection (**Supplemental Figure 3A**). Tissue samples were prepared from P84 male and female C57BL/6J mice and CoIP run in block design (**Supplemental Figure 3B-E**). Bilateral cerebral cortex and underlying structures from each brain were separated from the olfactory bulb at the optic chiasm and from tissue posterior to the cerebral cortex-midbrain junction (the cerebellum and majority of the brainstem). Brain tissue samples were homogenized in ice-cold Triton x-100 lysis buffer^17^ (1% Triton x-100, 5M NaCl, 25mM HEPES [ph 8],1mM EDTA [ph 8] in double distilled H2O) using a prechilled micro-pestle and passing homogenate through a 1mL syringe with a 23G needle ten times. Brain lysate was then incubated on ice for 10m to solubilize proteins and centrifuged for 20m at 14,000g at 4°C to remove the insoluble fraction. Supernatant was then transferred to a new tube and centrifuged for 10m at 12,000g at 4°C. Supernatant was RNAse treated on ice for 30m. Protein concentration was estimated using the BCA Protein Assay Kit described above (**Supplemental Figure 3B**). 50μL input samples were generated prior to supernatant combination with antibody-coupled beads. Antibody-coupled beads were prepared by incubating 3mg Dynabeads (Thermo Fisher Scientific #10018D) in 30μg mouse anti-Ca_V_β4 (Neuromab #75-054) overnight. Supernatant was precleared in 1.5mg 1xPBS pre-washed Dynabeads on a rotator for 1h at 4°C. Precleared supernatant was then incubated with antibody coupled beads (one “β4-IP” sample per mouse), or pre-washed Dynabeads beads not coupled to antibody (one “CN” per mouse)(total=16 samples overall) on rotator overnight at 4°C. β4-IP and CN samples were washed three times in ice-cold 1xPBS buffer and heated at 95°C for 5min in 25μL 1 x SDS and 10% mercaptoethanol in 4x protein sample loading buffer (LiCor #928-40004) to elute protein from beads.

#### Liquid Chromatography Mass Spectrometry (LC-MS/MS)

1μL (1667fmol) ^13^C_6_^15^N_4_-L-Arginine^13^C_6_^15^N_2_-L-Lysine Stable Isotope Labeled (SIL) β4 protein standard (Origene #PH310440) was added to each sample (total=16 samples overall) (**Supplemental Figure 3C**). Samples were ordered and prepared in a randomized balanced block design. Samples were digested with trypsin using the high-yield S-Trap™ Micro Spin columns (Protifi #C02-micro-80) following manufacture protocol and desalted using 0.6μL C18 resin ZipTips (Millipore Sigma #ZTC18S096). A pooled instrument control (PIC) was made by combining 1μL from each sample (**Supplemental Figure 3D**). 2μL (~1μg) each sample plus three 2μL aliquots of the PIC were resolved on an EASY C18 column (1.7μm, 2.1×50cm) at 300 nL/min with an UltiMate™ 3000 RSLCnano HPLC system over a 90-min gradient and analyzed on an Orbitrap Eclipse™ Tribrid™ MS (Thermo Fisher Scientific) operated in MS/MS after a final sample reordering (**Supplemental Figure 3E**). Peptide/protein identification, quantification and initial peptide peak area normalization were performed in Skyline Software^51^.

### Statistics

#### Quantitative Microscopy

Statistical tests were performed at two levels: 1) mouse level where n=16 mice and 2) neuron level where n=170 neurons in SPSS software (IBM, Armonk, NY). Shapiro-Wilk test was used to confirm normality. For statistical assessment at mouse level, an ANOVA (α=0.05) was used to test the main effects of group, sex, and group by sex interaction on mean DSD. These data were further examined at the neuron level in a separate statistical test. First, potential effects of mean distance of dendritic branch from soma, mean number of GFP(+)mCherry(+) cells in ROI and region were evaluated for any association with DSD. None were significantly associated with DSD and were thus not included in the final model. The final model to test spine density was an ANOVA (α=0.05) to test the main effects of group, sex, and group by sex interaction on DSD. The final model to test significance of impact of β4OE on density of spines based on volume was an ANOVA (α=0.05) which tested main effects of group, sex and group by sex interaction on DSD of each individual spine mask size bin. MANOVA (α=0.05) with Bonferroni correction was the final model used to detect significant differences in mean densities among dendritic protrusion types; main effects of group, sex and group by sex interaction were again tested with eight dependent variables: mean short stubby, long stubby, thin, short mushroom, long mushroom, branched, atypical and filopodia densities.

#### Western Blot

Mean optical density for β4OE, CN and AAV Naïve were calculated by averaging β-tubulin normalized technical replicates. Mean optical density β4OE versus AAV Naïve was evaluated via t-test, as was mean optical density of β4OE versus CN. ANOVA was used to detect differences in mean optical density of groups of male and female wildtype mice. Mean optical density in the following cases was calculated by averaging β-tubulin normalized optical density of β protein technical replicates for each mouse and then creating group means. For each β protein assessment, blot significantly impacted the primary outcome measure: mean β/mean β-tubulin, therefore, blot was included as a covariate in the statistical analysis for each β protein assessed (β1, β3, and β4).

#### CoIP-MS

β4 interactome data was analyzed by our group as previously described^41^. Briefly, within each sample peptide peak areas were exported from Skyline Software. β4 peptide levels were then normalized to SIL β4 peptide levels to control for differences in efficiency of the IP. Next, within each CoIP, peptide peak areas were divided by the normalized β4 peptide levels to generate normalized peak area ratios. Mean overall (includes samples from male and female mice) β4-IP peak area and mean overall CN peak area ratios were then calculated. Next, log2 fold change values were calculated using the overall means to detect shifts in distributions of β4-IP enriched protein levels, relative to CN. Peptides with CV<0.3 were compared using paired Student’s t-tests of log2-transformed peak ratios. Significantly altered peptides were defined as those with p value less than 0.05, p value greater than critical value (calculated using false discovery rate at 0.05) and log2 fold change>1 (peptides enriched >100% in males) or log2 fold change<−1 (peptides enriched >100% in females).

## RESULTS

### β4OE reduced DSD of L5 pyramidal cells in sensory cortex

A brief overview of mouse and cell numbers (see **Supplemental Figure 1**): Twenty-four P0-P2 C57BL/6J mouse pups were exposed to AAV injectate (12 to injectate A) using the BReVI procedure. Two males and two females perished prior to P84 due to either dam neglect (n=3) or hydrocephaly (n=1). Twenty mice were euthanized on P84: 4 male CN, 6 male β4OE, 4 female CN, and 6 female β4OE mice. A sparse labeling approach was utilized to avoid over-sampling spines, with the tradeoff that 4 (1 male CN, 2 male β4OE and 1 female CN) of the 20 total mice euthanized were excluded from this assessment prior to image analysis, due to paucity of L5 GFP(+)mCherry(+) cells visible at 10x. Sixteen total mice were assessed and for each mouse all L5 GFP(+)mCherry(+) cells visible at 10x were imaged: 3 male CN (n=42 total cells), 4 male β4OE (n=28 cells), 3 female CN (n=30 cells) and 6 female β4OE mice (n=70 cells). We conducted analyses of DSD at the level of mouse and at the level of neuron, focusing on the latter, since group sizes with n=cell were better balanced, providing the ability to test sex effects.

At the mouse level, β4OE significantly reduced mean DSD on minor basal branches of L5 pyramidal cells in sensory cortex (V1, V2, A1, A2, and TeA)(F=9.249, DF=3, p=0.01, **Figure 1B**). At the neuron level, β4OE significantly reduced DSD (F=30.922, DF=3, p<0.001, **Figure 1C**). Although the main effect of sex on DSD was not significant, there was a significant group by sex interaction (F=16.372, DF=3, p<0.001). In secondary analyses that considered DSD of neurons from CN versus β4OE in the two sexes independently, β4OE significantly reduced DSD in female (F=61.899, DF=1, p<0.001) but not in male mice relative to CN (**Figure 1D-E**).

### β4OE-mediated DSD reduction in female mice is independent of estrous stage

Female CN and β4OE mice assessed in this study were analyzed in the following estrous stages: proestrus, estrus and metestrus/diestrus. Neuron DSD of female mice is displayed broken down by estrous stage in **Figure 2A**. Because our determination of estrous stage was post-hoc, resulting in a non-random distribution of β4OE across estrous groups, we undertook several further analyses to elucidate whether β4OE reduced DSD in female mice, or if the apparent reduction was a spurious association due to confounding by estrous stage. Metestrus/diestrus was the only stage with neurons from both CN and β4OE mice. DSD was significantly reduced in female β4OE mice in metestrus/diestrus alone (F=6.190, DF=1, p=0.017, **Figure 2B**). Next, we randomly selected three β4OE female mice (Ms10-87, Ms5-10 and Ms16-169) and conducted within mouse assessments. DSD of β4OE+ neurons was significantly decreased compared to DSD of β4OE-internal control neurons overall (F=103.646, DF=1, p<0.001), regardless of estrous stage; there was a significant interaction between estrous stage and condition (β4OE+ versus β4OE-) (F=8.105, DF=1, p=0.006), however main effect of estrous stage was not significant (F=0.985, DF=1, p=0.324). DSD of β4OE+ neurons was significantly lower in each mouse: Ms10-87 was in estrus (F=67.688, DF=1, p<0.001), and Ms5-10 (F=9.502, DF=1, p=0.004) and Ms16-169 (F=126.518, DF=1, p<0.001) were in metestrus/diestrus (**Figure 2C**). Additionally, DSD of β4OE-neurons across the three β4OE mice assessed in **Figure 2C** did not significantly differ based on estrous stage (F=2.108, DF=1, p=0.154)(**Figure 2D**).

### β4OE selectively reduced density of spines with small volumes in female mice

Because spine density reductions due to β4OE were limited to females, our subsequent spine analyses focused on female mice. Mean DSD of small < 0.1μm^3^ spines was significantly decreased in neurons from female β4OE mice, relative to female CN (F=5.276, DF=1, p=0.035). Mean DSD of larger spine objects > 0.1μm^3^ did not significantly differ based on group in female mice (**Figure 3A**). β4OE significantly decreased mean DSD of small spines 0.1μm^3^ - 0.2μm^3^ in volume (F=6.748, DF=1, p=0.032) in female β4OE mice in metestrus/diestrus alone (**Figure 3B**). The effect of reduced small spine objects was not observed in male mice (**Supplemental Figure 4A**).

### β4OE decreased density of dendritic protrusions in female mice

β4OE significantly decreased mean density of four dendritic spine morphologic categories: short stubby, long stubby, short mushroom and long mushroom in female mice (**Figure 3C**). These four categories comprised the majority (95.19%) of all dendritic protrusions observed in females at P84. Focusing exclusively on females in metestrus/diestrus, β4OE significantly decreased mean density of short mushroom spines (F=20.160, DF=1, p<0.001)(**Figure 3D**). In contrast to effects observed in females, alterations to the spine morphologic categories of males were subtle (**Supplemental Figure 4B**).

### β subunit protein levels varied based on age and sex in C57BL/6J mice

β4 levels in A1 were significantly lower at P84, compared to P28, in male and female mice (F=9.635, DF=3, p=0.013, **Figure 1B**), with no significant sex or age by sex interaction. β1 levels were significantly lower in P84, relative to P28 male and female mice (F=21.499, DF=3, p=0.001). There was also a main effect of sex (F=6.944, DF=3, p=0.027) and a significant age by sex interaction (F=6.835, DF=3, p=0.028). β1 levels significantly differed in males versus females at P28 (F=12.138, DF=1, p=0.040) but not at the P84 timepoint (F=0.016, DF=1, p=0.906). Neither age nor sex significantly impacted levels of β3. Similarly, there was not a significant sex by age interaction.

### β1b significantly enriched in the β4 interactome of male C57BL/6J mice

The validity of the CoIP LC-MS approach was confirmed by identifying the 13023 peptides that comprise the β4 interactome in P84 mouse brain homogenate and verifying expected peptide/proteins consistent with previous reports of proteins associated with or predicted to be associated with the β subunit or β4 specifically^2^(**Supplemental Table 1**). Of the expected peptide/protein identifications, most notably, the VGCC subunits Ca_V_2.1, Ca_V_2.2, Ca_V_2.3 and α_2_δ1 were included in the list of protein/peptides detected using CoIP LC-MS in the current study.

Among the 13023 peptides in the β4 interactome (**Figure 5A**), eighteen peptides were significantly enriched in male mice (log2 fold change greater than 1 and p value less than 0.05). Twenty-four peptides in the β4 interactome were significantly enriched in females (log2 fold change less than −1 and p value less than 0.05, **Figure 5B**). Of particular interest was the peptide TMATAALAASPAPVSNLQGPYLASGDQPLDR, which was significantly enriched in the β4 interactome of male relative to female P84 mice (log2 fold change=3.9357, p=0.0201). This peptide is contained in the auxiliary VGCC subunit β1b. Amino acid sequence of this peptide was aligned to the amino acid sequences of each of the four β1 isoforms and revealed this peptide is exclusively found in the β1 isoform that is expressed in brain, β1b.

**Figure 4.**
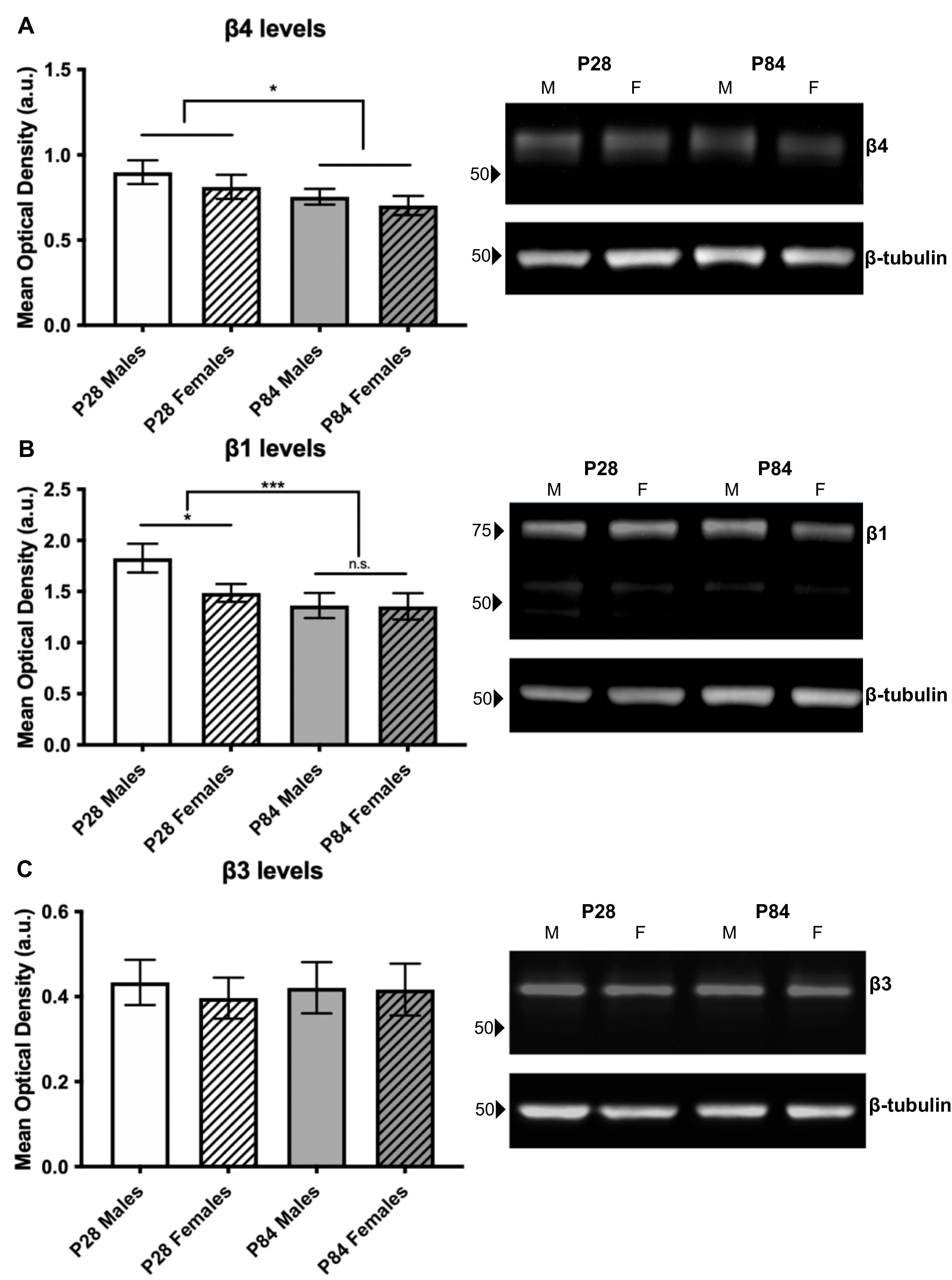
β subunit protein levels. **A)** (Left) Mean optical density (a.u.) of β4 was significantly lower in P84, relative to P28 male and female mice (F=9.635, DF=3, p=0.013), with no significant sex or age by sex interactions. Error bars = SEM. (Right) Representative blot showing β4 bands at ~51-55kD. **B)** (Left) Mean optical density (a.u.) of β1 was significantly lower in P84, relative to P28 male and female mice (F=21.499, DF=3, p=0.001). There was also a main effect of sex (F=6.944, DF=3, p=0.027) and a significant age by sex interaction (F=6.835, DF=3, p=0.028). Main effect of sex was significant at P28 (F=12.138, DF=1, p=0.040) but not P84 (F=0.016, DF=1, p=0.906). Error bars = SEM. (Right) Representative blot showing β1 bands at ~50-80kD. **C)** (Left) Neither age nor sex significantly impacted mean optical density (a.u.) of β3 and there was not a significant sex by age interaction. Error bars = SEM. (Right) Representative blot showing β3 bands at ~55kD (molecular weight of the predominant β3 isoform).

**Figure 5.**
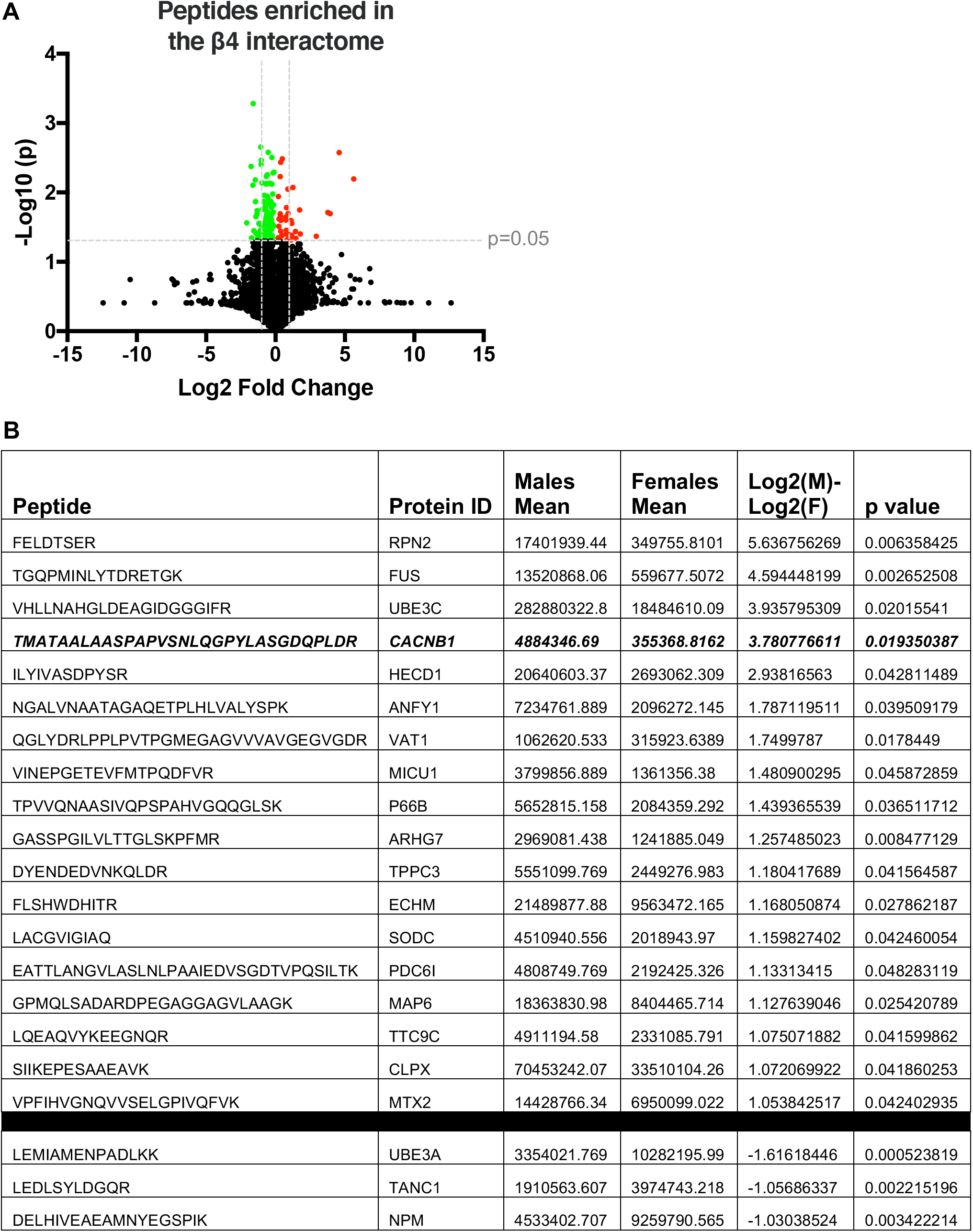
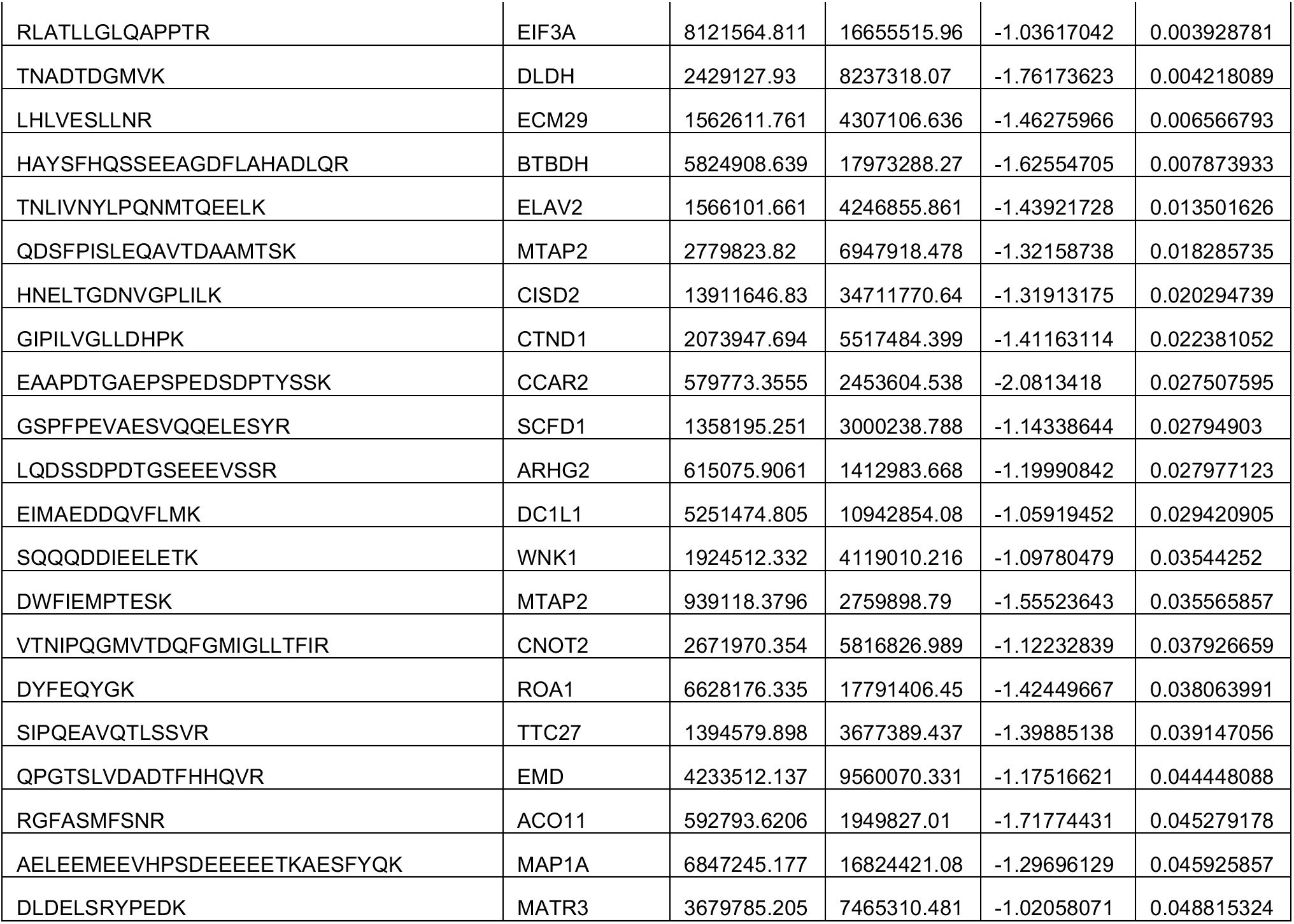
Sex differences in the β4 interactome of male relative to female P84 mice. **A)** Volcano plot showing 13023 β4-IP enriched peptides that make up the β4 interactome. Enriched peptides with p<0.05 are represented either in green (enriched in females) or red (enriched in males). Grey vertical lines (*L-R*) located at log2fold change = −1, 1. **B)** Table with the 18 significantly enriched peptides in males (with log2 fold change>1) and 24 significantly enriched peptides in females (with log2 fold change<−1). TMATAALAASPAPVSNLQGPYLASGDQPLDR (bolded, italicized) is the primary peptide of interest in these results; this peptide is significantly enriched in the β4 interactome of male relative to female P84 wildtype mice and is contained in the b isoform of the β1 subunit of voltage-gated calcium channels.

## DISCUSSION

In the current study, we tested the hypothesis that overexpressing CACNB4 in the developing brain decreases DSD in adult mice, mirroring the spine loss anatomical phenotype observed in A1 in postmortem schizophrenia^30,33,34,52^. Although β4OE did indeed significantly reduce DSD of L5 pyramidal cells in sensory cortex of female mice, this was not the case for male mice. We evaluated whether this sex difference was driven by estrous stage rather than β4OE itself by examining DSD of CN and β4OE females during all estrous stages, during metestrus/diestrus alone, and comparing DSD of β4OE+ to DSD of β4OE- (internal control) pyramidal cells in female β4OE mice. The results from these assessments provided convincing evidence that the spine loss observed in β4OE female mice was not an artifact of estrous stage differences between groups. Spine volume assessment revealed β4OE-driven spine loss in female mice was selective for loss of spines of small volumes, recapitulating the small spine loss in A1 in schizophrenia^30,33,34,52^. Possibly underlying the sex-specific effect of β4OE, our analyses revealed that the β1 VGCC subunit expressed in brain, β1b, was significantly enriched in the β4 interactome of male compared to female adult C57Bl/6J mice.

Two methodological limitations encountered in these studies warrant discussion. First, we intentionally employed a sparse labeling approach to assess dendritic spines on pyramidal cells in order to: clearly visualize spines with optimal signal-to-noise characteristics and avoid over-sampling spine objects in our tissue. A minor downside of making this choice is that it necessitated expanding the ROI from A1, the brain region specified in our scientific premise, to other regions of sensory cortex (V1, V2, A1, A2, and TeA)^30,33,34,52^. However, we and others have shown that unlike in higher mammals^53–56^, spine density does not significantly differ based on region in the cerebral cortex of mice^57–61^. The major limitation of sparse labeling in the current study is that this led to an uneven number of mice per group. Since further reducing the size of the groups before image processing to equalize the numbers was not sensible, we attempted to mitigate this limitation using multiple approaches: including as many controls as possible, probing all variables identified as potential artifacts or confounds and reporting DSD at both mouse and neuron levels following rigorous statistical testing. A second limitation is that the P2A element included in the β4OE AAV we used was not 100% efficient resulting in the expression of a β4-GFP fusion protein in addition to non-native β4 and GFP in β4OE mice. One concern is that the fusion protein behaves differently than simple overexpression of non-native β4. However, overexpressing β4, which lacked a GFP tag, in primary cortical cultures, significantly decreased DSD in our previous study^34^, just as β4OE significantly reduced DSD in female mice in the current study.

Several influential light and electron microscopy studies have demonstrated that circulating hormone levels across the estrous cycle shape spine density in hippocampus of female rats^62^ with elevated levels of 17β-estradiol (E2) during proestrus associated with higher DSD^63–65^. Results from similar studies performed in rodent cortex, however, are mixed. Two out of three studies in rat cortex found no difference in DSD in proestrus compared to other stages^66–68^. Studies of mouse cortex indicate no change in DSD due to estrous stage: DSD of L5 pyramidal cells across estrous in somatosensory cortex was unchanged^62^, as was DSD of L5 pyramidal cells in frontal cortex in hormone-treated versus ovariectomized female mice^69^. Nevertheless, we explored the possibility that our finding of β4OE-mediated spine loss in female mice could be driven by estrous stage by evaluating vaginal cytology of each female mouse included in our spine study. All animals in the study were sacrificed on a single day, P84, since DSD is known to fluctuate over the course of neurodevelopment as a function of age, specifically in A1 and sensory cortex of mice^43,50^. Given that each estrous stage was not equally represented on P84 across groups by chance, we specifically compared DSD of female CN versus β4OE mice in metestrus/diestrus, the only estrous stage represented in both groups of mice. Narrowing our focus to metestrus/diestrus again revealed DSD was significantly reduced in female β4OE relative to CN mice, replicating at a smaller scale the overall pattern in females. Importantly, DSD of β4OE+ neurons was significantly decreased compared to DSD of internal control neurons in a β4OE mouse in estrus. The same finding was repeated in two β4OE mice in metestrus/diestrus. Further, DSD of internal control neurons did not significantly differ based on estrous stage in β4OE mice. These data collectively indicate that β4OE-mediated DSD reduction in female mice was not an artifact of estrous stage (**Figure 2B-D**). Moreover, our internal control assessment findings suggest β4OE exerts a cell-autonomous effect as DSD was not reduced in internal control neurons in β4OE mice.

The predominant roles of voltage-gated calcium channel β subunits are to forward traffick the α1 subunit of VGCCs toward the plasma membrane and regulate the biophysical properties of the channel^19–22^. In heterologous expression systems, VGCC calcium currents substantially increase when β subunits are coexpressed with α1 subunits, compared to expression of α1 subunits alone^2^,^3^. Further, all β subunits readily homooligomerize and this oligomerization selectively increases VGCC calcium current density^70–72^. Therefore, it seems likely that β4OE increased calcium current density through VGCCs in the current study. Post-synaptic calcium spikes are compulsory for initiating signaling cascades that shape synaptic plasticity induction and actin cytoskeleton remodeling^23^,^26^,^27^. If β4OE mediates calcium entry into spines at levels that exceed the regulatory capacity of these spines, synaptotoxicity and subsequent spine loss may result. Synaptotoxicity, a feature of early pathogenesis, is well described in Alzheimer’s disease. Aβ oligomer-induced synaptotoxicity is characterized by reduced density of excitatory synapses^73–76^; several underlying molecular mechanisms of elevated cytosolic calcium at the plasma membrane have been suggested. Aβ aggregates elicit calcium dyshomeostasis, resulting in increased cytosolic calcium in neurons, including at synaptic sites^77^. Soluble Aβ oligomers increase Ca_V_1-mediated calcium entry^78–80^, decrease cytosolic calcium efflux via plasma membrane calcium ATPase and/or Na^+^/Ca^2+^ exchanger inhibition, although results concerning exchangers are mixed^81–84^, and perforate cell membranes, allowing calcium to flow unhindered down a steep concentration gradient^85^,^86^. 24h treatment with 25μM Aβ increased protein and surface expression of Ca_V_1.2 and Ca_V_1.3 channels in hippocampal neurons^78^,^79^. 1h treatment with 1μM Aβ increased Ca_V_1 currents in rat cortical synaptosomes, whereas 16h Aβ treatment (1μM) increased Ca_V_1 calcium currents in rat cortical neurons^80^. In neuroblast cells intracellular Aβ expression increased cytosolic calcium and Ca_V_1 blockers prevented amyloid beta protein precursor C-terminal fragment-mediated neurotoxicity^87^. Finally, emerging research suggests synptotoxicity and subsequent dendritic spine loss in Parkinson’s disease may result from significantly increased Ca_V_1.3-mediated calcium influx following dopamine depletion^88,89^. Although synaptotoxicity is typically discussed in the neurodegeneration literature, the data presented and discussed herein may provide evidence that a similar molecular mechanism may underlie β4OE-mediated spine loss resported in this and our previous study^34^.

Could there be a potential explanation for why spines of small volumes are particularly susceptible to the β4OE manipulation? In the current study, spines of small volume (<0.3μm^3^) made up the majority (68.96%) of all spines observed in P84 mice. The actin cytoskeleton of small spines, many of which are transient and/or lacking a mature postsynaptic density, are particularly vulnerable to the destabilizing effects of altered calcium transients whether through activity-dependent synaptic plasticity or pathological mechanisms^90–92^. Recent modeling studies shed light on these observations, finding spine head volume to be inversely proportional to intracellular calcium content, due to calcium efflux and buffering source variables in spines with large surface area. Modeled spine volume was both inversely proportional to peak Ca^2+^ and area under the curve of Ca^2+^ ions at 300ms post-stimulus regardless of spine shape^93^. If β4OE does indeed increase calcium transients into spines of β4OE mice, the smallest spines would be the most susceptible to high calcium concentrations with the potential to induce synaptotoxicity and spine loss.

We observed significant small spine loss only in female β4OE mice, leading us to evaluate whether the interactome of β4 differed between males and females. Of note, we found enrichment of β1b, the β1 isoform expressed in brain, in the β4 interactome of male mice, suggesting that β1b might mitigate the effects of β4OE in males. Although the presence of β1b in the β4 interactome does not specifically indicate that the two proteins are directly interacting and it is currently unknown if β1b and β4 oligomerize, we cannot rule out this possibility. Heterooligomers comprising β1 and β3 have been previously reported^70^. If β4 and β1b do interact in brain, association is likely to take place, at least in part, at residues in the guanylate kinase subdomain of the membrane-associated guanylate kinase-like (MAGUK) domain of these proteins^70^. β2d-β3 oligomers alter the biophysical properties of Ca_V_1 shifting the peak I-V curve by 20mV in the depolarized direction compared to β2d or β3 monomer expression alone, with a slight but non-significant reduction in inactivation rate^70^. A sensible potential implication of β1b and β4 heterooligomerization is to behave like other β heterooligomers by shifting the peak I-V curve of Ca_V_1 channels to depolarized voltages, particuarly if β heterooligomers docked on α1 subunits do not elicit additional significant changes to channel gating. Significantly shifting peak I-V curve in the depolarized direction would be expected to decrease VGCC-mediated inward flow of calcium, an outcome that has the potential to protect against synaptotoxicity and spine loss.

Neither β1b protein nor β4 protein levels (predominant neuronal isoforms) were significantly different based on sex in P84 mouse brain, at least in A1 (**Figure 4A-B**). Although there was a significant effect of sex on β1b protein levels overall, and significant age by sex interaction, both were driven by the significant difference in β1b protein levels in males versus females at the P28 timepoint. Therefore, a sensible explanation for why β1b is significantly enriched in the β4 interactome of males *relative* to females (**Figure 5A-B**) is due to differential targeting/subcellular localization of β subunits in these animals based on sex. We are not aware of any studies that report or investigate mechanisms for differential targeting or subcellular localization of β VGCC subunits in neurons based on sex. This is not surprising as investigating sex differences has not been a widespread priority in biomedical research until recently^95^. However, newly published data reveal sex-dependent regulation of VGCC function of Ca_V_1 channels in dopamine neurons that innervate the striatum^96^, sex differences in Ca_V_1 function have also been documented in cardiac tissue^97,98^, and we expect future studies will discover additional sex divergence in the regulation, function and subcellular localization of primary and auxiliary VGCC subunits.

Our findings on one hand extend our previous *in vitro* finding of CACNB4 overexpression-mediated loss of small dendritic spines to an intact, developed mammalian system. They further reveal nuances including two notable sex differences: β4OE-mediated spine loss was significant in female but not male mice and the β1 VGCC subunit was significantly enriched in the β4 VGCC subunit interactome of male relative to female C57BL/6J mice. The current studies therefore help expand our understanding about mechanisms leading to structural spine plasticity generally, emphasizing the need to study both males and females^95^. Although our spine findings do not fully recapitulate the spine phenotype in schizophrenia, in which small spine loss in A1 extends to both sexes ^33,34^, perhaps this divergence is unsurprising. Schizophrenia is a polygenic disorder with hundreds of genetic variants and additional environmental exposures associated with risk^29,99,100^. Postmortem human studies demonstrating small spine loss include individuals who develop small spine loss via many different molecular mechanisms. Although our spine findings do not translate to the schizophrenia population as a whole, we cannot rule out the possibility that sex differences in DSD may be present in pairs of schizophrenia subjects with shared genetic liability, for example, in individuals that harbor variants that fundamentally alter post-synaptic calcium signaling.

## REFERENCES

1 Zamponi, G. W. Voltage-gated calcium channels. (Springer, (2005).

2 Buraei, Z. & Yang, J. The β subunit of voltage-gated Ca 2+ channels. Physiological Reviews 90, 1461–1506 (2010).

3 Buraei, Z. & Yang, J. Structure and function of the β subunit of voltage-gated Ca2+ channels. Biochimica Et Biophysica Acta (BBA)-Biomembranes 1828, 1530–1540 (2013).

4 Dolphin, A. C. Voltage-gated calcium channels and their auxiliary subunits: physiology and pathophysiology and pharmacology. The Journal of Physiology 594, 5369–5390 (2016).

5 Castellano, A. & Perez-Reyes, E. Molecular diversity of Ca2+ channel β subunits. Biochemical Society Transations 22, 483–488 (1994).

6 Schlick, B., Flucher, B. & Obermair, G. Voltage-activated calcium channel expression profiles in mouse brain and cultured hippocampal neurons. Neuroscience 167, 786–798 (2010).

7 Yeon, J.-H., Park, C.-G., Hille, B. & Suh, B.-C. Translocatable voltage-gated Ca2+ channel β subunits in α1–β complexes reveal competitive replacement yet no spontaneous dissociation. Proceedings of the National Academy of Sciences 115, E9934–E9943 (2018).

8 Burgess, D. L. et al. β subunit reshuffling modifies N-and P/Q-type Ca 2+ channel subunit compositions in lethargic mouse brain. Molecular and Cellular Neuroscience 13, 293–311 (1999).

9 Burgess, D. L., Jones, J. M., Meisler, M. H. & Noebels, J. L. Mutation of the Ca2+ channel β subunit gene Cchb4 is associated with ataxia and seizures in the lethargic (lh) mouse. Cell 88, 385–392 (1997).

10 Etemad, S. et al. Differential neuronal targeting of a new and two known calcium channel β4 subunit splice variants correlates with their regulation of gene expression. Journal of Neuroscience 34, 1446–1461 (2014).

11 Ludwig, A., Flockerzi, V. & Hofmann, F. Regional expression and cellular localization of the α1 and β subunit of high voltage-activated calcium channels in rat brain. Journal of Neuroscience 17, 1339–1349 (1997).

12 Ferrándiz-Huertas, C., Gil-Mínguez, M. & Luján, R. Regional expression and subcellular localization of the voltage-gated calcium channel β subunits in the developing mouse brain. Journal of Neurochemistry 122, 1095–1107 (2012).

13 Tanaka, O., Sakagami, H. & Kondo, H. Localization of mRNAs of voltage-dependent Ca2+-channels: four subtypes of α1-and β-subunits in developing and mature rat brain. Molecular Brain Research 30, 1–16 (1995).

14 Wittemann, S., Mark, M. D., Rettig, J. & Herlitze, S. Synaptic localization and presynaptic function of calcium channel β4-subunits in cultured hippocampal neurons. Journal of Biological Chemistry 275, 37807–37814 (2000).

15 Scott, V. E. et al. Beta subunit heterogeneity in N-type Ca2+ channels. Journal of Biological Chemistry 271, 3207–3212 (1996).

16 McEnery, M. W. et al. Differential expression and association of calcium channel subunits in development and disease. Journal of Bioenergetics and Biomembranes 30, 409–418 (1998).

17 Klemmer, P., Smit, A. B. & Li, K. W. Proteomics analysis of immuno-precipitated synaptic protein complexes. Journal of Proteomics 72, 82–90 (2009).

18 Subramanyam, P. et al. Activity and calcium regulate nuclear targeting of the calcium channel beta4b subunit in nerve and muscle cells. Channels 3, 343–355 (2009).

19 Gonzalez-Gutierrez, G., Miranda-Laferte, E., Naranjo, D., Hidalgo, P. & Neely, A. Mutations of nonconserved residues within the calcium channel α1-interaction domain inhibit β-subunit potentiation. Journal of General Physiology 132, 383–395 (2008).

20 Maltez, J. M., Nunziato, D. A., Kim, J. & Pitt, G. S. Essential Ca v β modulatory properties are AID-independent. Nature Structural & Molecular Biology 12, 372–377 (2005).

21 Josephson, I. R. & Varadi, G. The beta subunit increases Ca2+ currents and gating charge movements of human cardiac L-type Ca2+ channels. Biophysical Journal 70, 1285–1293 (1996).

22 Jones, L. P., Wei, S.-k. & Yue, D. T. Mechanism of auxiliary subunit modulation of neuronal α1E calcium channels. Journal of General Physiology 112, 125–143 (1998).

23 Higley, M. J. & Sabatini, B. L. Calcium signaling in dendritic spines. Cold Spring Harbor Perspectives in Biology 4, a005686 (2012).

24 Sabatini, B. L. & Svoboda, K. Analysis of calcium channels in single spines using optical fluctuation analysis. Nature 408, 589–593 (2000).

25 Yasuda, R., Sabatini, B. L. & Svoboda, K. Plasticity of calcium channels in dendritic spines. Nature Neuroscience 6, 948–955 (2003).

26 Yuste, R., Majewska, A. & Holthoff, K. From form to function: calcium compartmentalization in dendritic spines. Nature Neuroscience 3, 653–659 (2000).

27 Kasai, H., Matsuzaki, M., Noguchi, J., Yasumatsu, N. & Nakahara, H. Structure– stability–function relationships of dendritic spines. Trends in Neurosciences 26, 360–368 (2003).

28 Ripke, S. et al. Genome-wide association analysis identifies 13 new risk loci for schizophrenia. Nature Genetics 45, 1150–1159 (2013).

29 Ripke, S. et al. Biological insights from 108 schizophrenia-associated genetic loci. Nature 511, 421–427 (2014).

30 Sweet, R. A., Henteleff, R. A., Zhang, W., Sampson, A. R. & Lewis, D. A. Reduced dendritic spine density in auditory cortex of subjects with schizophrenia. Neuropsychopharmacology 34, 374–389 (2009).

31 Moyer, C. E., Shelton, M. A. & Sweet, R. A. Dendritic spine alterations in schizophrenia. Neuroscience Letters 601, 46–53 (2015).

32 Shelton, M. A. et al. Loss of microtubule-associated protein 2 immunoreactivity linked to dendritic spine loss in schizophrenia. Biological Psychiatry 78, 374–385 (2015).

33 McKinney, B. C. et al. Density of small dendritic spines and microtubule-associated-protein-2 immunoreactivity in the primary auditory cortex of subjects with schizophrenia. Neuropsychopharmacology 44, 1055–1061 (2019).

34 MacDonald, M. L. et al. Selective loss of smaller spines in schizophrenia. American Journal of Psychiatry 174, 586–594 (2017).

35 Gholizadeh, S., Tharmalingam, S., MacAldaz, M. E. & Hampson, D. R. Transduction of the central nervous system after intracerebroventricular injection of adeno-associated viral vectors in neonatal and juvenile mice. Human Gene Therapy Methods 24, 205–213 (2013).

36 Stoica, L., Ahmed, S. S., Gao, G. & Esteves, M. S. AAV-mediated gene transfer to the mouse CNS. Current Protocols in Microbiology, Chapter 14, Unit 14D.15 (2013).

37 Cheetham, C. E. J., Grier, B. D. & Belluscio, L. Bulk regional viral injection in neonatal mice enables structural and functional interrogation of defined neuronal populations throughout targeted brain areas. Frontiers in Neural Circuits 9, doi:10.3389/fncir.2015.00072 (2015).

38 Phifer, C. B. & Terry, L. M. Use of hypothermia for general anesthesia in preweanling rodents. Journal of Physiology Behavior 38, 887–890 (1986).

39 McLean, A. C., Valenzuela, N., Fai, S. & Bennett, S. A. Performing vaginal lavage, crystal violet staining, and vaginal cytological evaluation for mouse estrous cycle staging identification. JoVE e4389, doi:10.3791/4389 (2012).

40 Byers, S. L., Wiles, M. V., Dunn, S. L. & Taft, R. A. Mouse estrous cycle identification tool and images. PLoS One 7, e35538, doi:10.1371/journal.pone.0035538 (2012).

41 Grubisha, M. J. et al. MAP2 is Hyperphosphorylated in Schizophrenia and Alters its Function. bioRxiv, 683912 (2019).

42 Franklin, K. B. J. & Paxinos, G. The mouse brain in stereotaxic coordinates. (Gulf Professional Publishing, 2004).

43 Parker, E. M., Kindja, N. L., Cheetham, C. E. J. & Sweet, R. A. Sex differences in dendritic spine density and morphology in auditory and visual cortices in adolescence and adulthood. Scientific Reports 10, 1–11, doi:10.1038/s41598-020-65942-w (2020).

44 Bopp, R., Holler-Rickauer, S., Martin, K. A. C. & Schuhknecht, G. F. P. An ultrastructural study of the thalamic input to layer 4 of primary motor and primary somatosensory cortex in the mouse. Journal of Neuroscience 37, 2435–2448 (2017).

45 Zhang, W., Peterson, M., Beyer, B., Frankel, W. N. & Zhang, Z.-w. Loss of MeCP2 from forebrain excitatory neurons leads to cortical hyperexcitation and seizures. Journal of Neuroscience 34, 2754–2763 (2014).

46 Li, H.-S. et al. Inactivation of Numb and Numblike in embryonic dorsal forebrain impairs neurogenesis and disrupts cortical morphogenesis. Neuron 40, 1105–1118 (2003).

47 Arellano, J. I., Benavides-Piccione, R., DeFelipe, J. & Yuste, R. Ultrastructure of dendritic spines: correlation between synaptic and spine morphologies. Frontiers in Neuroscience 1, 131–143 (2007).

48 Risher, W. C., Ustunkaya, T., Alvarado, J. S. & Eroglu, C. Rapid Golgi analysis method for efficient and unbiased classification of dendritic spines. PLoS One 9, e107591, doi:10.1371/journal.pone.0107591 (2014).

49 Caligioni, C. S. Assessing reproductive status/stages in mice. Current Protocols in Neuroscience, Appendix 4 48, Appendix–4I (2009).

50 Moyer, C. E. et al. Developmental trajectories of auditory cortex synaptic structures and gap-prepulse inhibition of acoustic startle between early adolescence and young adulthood in mice. Cerebral Cortex 26, 2115–2126 (2015).

51 MacLean, B. et al. Skyline: an open source document editor for creating and analyzing targeted proteomics experiments. Bioinformatics 26, 966–968 (2010).

52 Shelton, M. A. et al. Loss of microtubule-associated protein 2 immunoreactivity linked to dendritic spine loss in schizophrenia. Biological psychiatry 78, 374–385 (2015).

53 Amatrudo, J. M. et al. Influence of highly distinctive structural properties on the excitability of pyramidal neurons in monkey visual and prefrontal cortices. Journal of Neuroscience 32, 13644–13660 (2012).

54 Gilman, J. P., Medalla, M. & Luebke, J. I. Area-specific features of pyramidal neurons—a comparative study in mouse and rhesus monkey. Cerebral Cortex 27, 2078–2094 (2016).

55 Jacobs, B. et al. Regional Dendritic and Spine Variation in Human Cerebral Cortex: a Quantitative Golgi Study. Cerebral Cortex 11, 558–571 (2001).

56 Clemo, R. H. & Meredith, A. M. Dendritic Spine Density in Multisensory Versus Primary Sensory Cortex. Synapse 66, 714–724 (2012).

57 Harris, K. D. & Shepherd, G. M. G. The neocortical circuit: themes and variations Nature Neuroscience 18, 170–181 (2015).

58 Arellano, J., Benavides-Piccione, R., DeFelipe, J. & Yuste, R. Ultrastructure of dendritic spines: correlation between synaptic and spine morphologies. Frontiers in Neuroscience 1, doi:10.3389/neuro.01.1.1.010.2007 (2007).

59 Benavides-Piccione, R., Ballesteros-Yáñez, I., DeFelipe, J. & Yuste, R. Cortical area and species differences in dendritic spine morphology. Journal of Neurocytology 31, 337–346 (2002).

60 Hsu, A., Luebke, J. I. & Medalla, M. Comparative ultrastructural features of excitatory synapses in the visual and frontal cortices of the adult mouse and monkey. Journal of Comparative Neurology 525, 2175–2191 (2017).

61 Luebke, J. I. Pyramidal neurons are not generalizable building blocks of cortical networks. Frontiers in Neuroanatomy 11, doi:10.3389/fnana.2017.00011 (2017).

62 Alexander, B. H. et al. Stable density and dynamics of dendritic spines of cortical neurons across the estrous cycle while expressing differential levels of sensory-evoked plasticity. Frontiers in Molecular Neuroscience 11, 83, doi:10.3389/fnmol.2018.00083 (2018).

63 Woolley, C. S., Gould, E., Frankfurt, M. & McEwen, B. S. Naturally occurring fluctuation in dendritic spine density on adult hippocampal pyramidal neurons. Journal of Neuroscience 10, 4035–4039 (1990).

64 Woolley, C. S. & McEwen, B. S. Estradiol mediates fluctuation in hippocampal synapse density during the estrous cycle in the adult rat. Journal of Neuroscience 12, 2549–2554 (1992).

65 Kato, A. et al. Female hippocampal estrogens have a significant correlation with cyclic fluctuation of hippocampal spines. Frontiers in Neural Circuits 7, 149, doi:10.3389/fncir.2013.00149 (2013).

66 Chen, J.-R. et al. Gonadal hormones modulate the dendritic spine densities of primary cortical pyramidal neurons in adult female rat. Cerebral Cortex 19, 2719–2727 (2009).

67 Markham, J. A. & Juraska, J. M. Aging and sex influence the anatomy of the rat anterior cingulate cortex. Neurobiology of Aging 23, 579–588 (2002).

68 Prange-Kiel, J. et al. Gonadotropin-releasing hormone regulates spine density via its regulatory role in hippocampal estrogen synthesis. Journal of Cell Biology 180, 417–426 (2008).

69 Boivin, J. R., Piekarski, D. J., Thomas, A. W. & Wilbrecht, L. Adolescent pruning and stabilization of dendritic spines on cortical layer 5 pyramidal neurons do not depend on gonadal hormones. Developmental Cognitive Neuroscience 30, 100–107 (2018).

70 Lao, Q. Z., Kobrinsky, E., Liu, Z. & Soldatov, N. M. Oligomerization of Cavβ subunits is an essential correlate of Ca2+ channel activity. The FASEB Journal 24, 5013–5023 (2010).

71 Hullin, R. et al. Increased expression of the auxiliary β2-subunit of ventricular L-type Ca2+ channels leads to single-channel activity characteristic of heart failure. PLoS One 2, e292 (2007).

72 Chen, X. et al. Ca2+ influx–induced sarcoplasmic reticulum Ca2+ overload causes mitochondrial-dependent apoptosis in ventricular myocytes. Circulation research 97, 1009–1017 (2005).

73 Rush, T. et al. Synaptotoxicity in Alzheimer’s disease involved a dysregulation of actin cytoskeleton dynamics through cofilin 1 phosphorylation. Journal of Neuroscience 38, 10349–10361 (2018).

74 Lacor, P. N. et al. Synaptic targeting by Alzheimer’s-related amyloid β oligomers. Journal of Neuroscience 24, 10191–10200 (2004).

75 Lacor, P. N. et al. Aβ oligomer-induced aberrations in synapse composition, shape, and density provide a molecular basis for loss of connectivity in Alzheimer’s disease. Journal of Neuroscience 27, 796–807 (2007).

76 Kervern, M. et al. Selective impairment of some forms of synaptic plasticity by oligomeric amyloid-β peptide in the mouse hippocampus: implication of extrasynaptic NMDA receptors. Journal Of Alzheimer’s disease 32, 183–196 (2012).

77 Green, K. N. Calcium in the initiation, progression and as an effector of Alzheimer’s disease pathology. Journal of cellular and molecular medicine 13, 2787–2799 (2009).

78 Min, D. et al. The alterations of Ca2+/calmodulin/CaMKII/CaV1.2 signaling in experimental models of Alzheimer’s disease and vascular dementia. Neuroscience letters 538, 60–65 (2013).

79 Kim, S. & Rhim, H. Effects of amyloid-β peptides on voltage-gated L-type Ca v1.2 and Ca v 1.3 Ca 2+ channels. Molecules and cells 32, 289–294 (2011).

80 Ueda, K., Shinohara, S., Yagami, T., Asakura, K. & Kawasaki, K. Amyloid β protein potentiates Ca2+ influx through L-type voltage-sensitive Ca2+ channels: a possible involvement of free radicals. Journal of neurochemistry 68, 265–271 (1997).

81 Mata, A. M. Functional interplay between plasma membrane Ca2+-ATPase, amyloid β-peptide and tau. Neuroscience letters 663, 55–59 (2018).

82 Kim, H.-S., Lee, J.-H. & Suh, Y.-H. C-terminal fragment of Alzheimer’s amyloid precursor protein inhibits sodium/calcium exchanger activity in SK-N-SH cell. Neuroreport 10, 113–116 (1999).

83 Colvin, R. A., Davis, N., Wu, A., Murphy, C. A. & Levengood, J. Studies of the mechanism underlying increased Na+/Ca2+ exchange activity in Alzheimer’s disease brain. Brain research 665, 192–200 (1994).

84 Moriguchi, S. et al. Reduced expression of Na+/Ca2+ exchangers is associated with cognitive deficits seen in Alzheimer’s disease model mice. Neuropharmacology 131, 291–303 (2018).

85 Sepulveda, F. J., Parodi, J., Peoples, R. W., Opazo, C. & Aguayo, L. G. Synaptotoxicity of Alzheimer beta amyloid can be explained by its membrane perforating property. PLoS One 5, e11820 (2010).

86 Pacheco, C. R. et al. Extracellular α-synuclein alters synaptic transmission in brain neurons by perforating the neuronal plasma membrane. Journal of neurochemistry 132, 731–741 (2015).

87 Anekonda, T. S. et al. L-type voltage-gated calcium channel blockade with isradipine as a therapeutic strategy for Alzheimer’s disease. Neurobiol. Dis. 41, 62–70 (2011).

88 Day, M. et al. Selective elimination of glutamatergic synapses on striatopallidal neurons in Parkinson disease models. Nature neuroscience 9, 251–259 (2006).

89 Oertner, T. G. & Matus, A. Calcium regulation of actin dynamics in dendritic spines. Cell Calcium 37, 477–482 (2005).

90 Murmu, R. P., Li, W., Holtmaat, A. & Li, J.-Y. Dendritic spine instability leads to progressive neocortical spine loss in a mouse model of Huntington’s disease. Journal of Neuroscience 33, 12997–13009 (2013).

91 Holtmaat, A. & Svoboda, K. Experience-dependent structural synaptic plasticity in the mammalian brain. Nature reviews. Neuroscience 10, 647 (2009).

92 Holtmaat, A. J. et al. Transient and persistent dendritic spines in the neocortex in vivo. Neuron 45, 279–291 (2005).

93 Bell, M., Bartol, T., Sejnowski, T. & Rangamani, P. Dendritic spine geometry and spine apparatus organization govern the spatiotemporal dynamics of calcium. Journal of General Physiology 151, 1017–1034 (2019).

94 Takahashi, E., Mitsuhiro, I. & Nagasu, T. Effect of Genetic Background on Cav2 Channel α1 and β Subunit Messenger RNA Expression in Cerebellum of N-Type Ca2+ Channel α1B Subunit-Deficient Mice. Comparative Medicine 54, 690–694 (2004).

95 Beery, A. K., Zucker, I.. Sex bias in neuroscience and biomedical research. Neurosci. Biobehav. Rev. 35, 565–572 (2011).

96 Brimblecombe, K. R. et al. L-type calcium channel contribution to striatal dopamine release is governed by calbindin-D28K, the dopamine transporter, D2-receptors, α2δ-subunits and sex differences. bioRxiv (2020).

97 Curl, C. L., Delbridge, L. M. & Wendt, I. R. Sex differences in cardiac muscle responsiveness to Ca2+ and L-type Ca2+ channel modulation. European journal of pharmacology 586, 288–292 (2008).

98 Prabhavathi, K., Selvi, K. T., Poornima, K. & Sarvanan, A. Role of biological sex in normal cardiac function and in its disease outcome–a review. Journal of clinical and diagnostic research: JCDR 8, BE01 (2014).

99 Purcell, S. M. et al. A polygenic burden of rare disruptive mutations in schizophrenia. Nature 506, 185–190 (2014).

100 Freedman, R. et al. Evidence for the multigenic inheritance of schizophrenia. Am. J. Med. Genet. 105, 794–800 (2001).

