## Supplement for "CACNB4 overexpression decreases dendritic spine density in sex-specific manner"

SUPPLEMENTAL FIGURES

**Supplemental Figure 1**

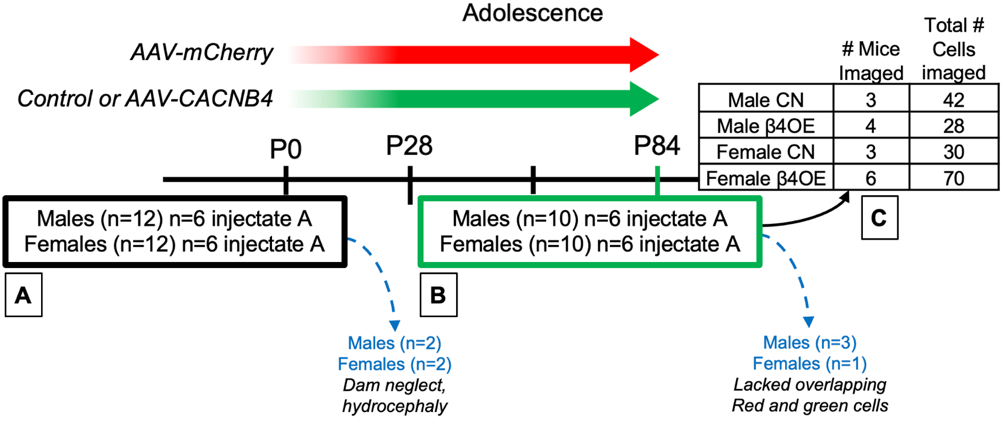

**Supplemental Figure 1** Dendritic spine study design and execution. **A)** Two versions of AAV injectate were prepared by adding AAV-mCherry to either control GFP AAV (to generate control [CN] mice) or AAV-CACNB4 (to generate β4 overexpression [β4OE] mice). To blind experimenter to group AAV solutions were randomly coded “A” and “B” prior to AAV injections. Twenty-four P0-P2 C57BL/6J mouse pups were exposed to AAV injectate (50% to injectate A) using the BReVI procedure. Two males and two females perished prior to P84 due to either dam neglect (n=3) or hydrocephaly (n=1). **B)** Stage of estrous cycle was estimated by evaluating vaginal cytology of P84 females on day of sacrifice. Twenty total mice were euthanized on P84: 4 male CN, 6 male β4OE, 4 female CN, and 6 female β4OE mice. Mice were anesthetized and transcardially perfused with ice-cold 1x PBS followed by 4% PFA. Brains were extracted, post-fixed in 4% PFA, moved to 18% sucrose and stored at -30°C in cryoprotectant until sectioning. **C)** Layer 5 (L5) mCherry fluorescent pyramidal cells with GFP somal labeling (GFP+mCherry+ cells) were selected for imaging. A sparse labeling approach was utilized to specifically avoid over-sampling spines, with the tradeoff that 4 (1 male CN, 2 male β4OE and 1 female CN) of the 20 total mice euthanized were excluded from this assessment prior to image analysis, due to paucity of GFP+mCherry+ cells visible at 10x. Sixteen total mice were assessed and for each mouse all GFP+mCherry+ cells visible at 10x were imaged: 3 male CN (n=42 total cells), 4 male β4OE (n=28 cells), 3 female CN (n=30 cells) and 6 female β4OE mice (n=70 cells).

**Supplemental Figure 2**

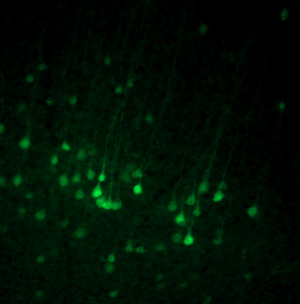

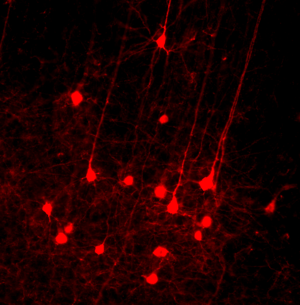

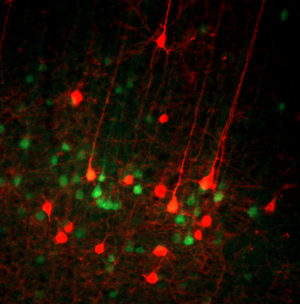

50μm

**GFP**

**mCherry**

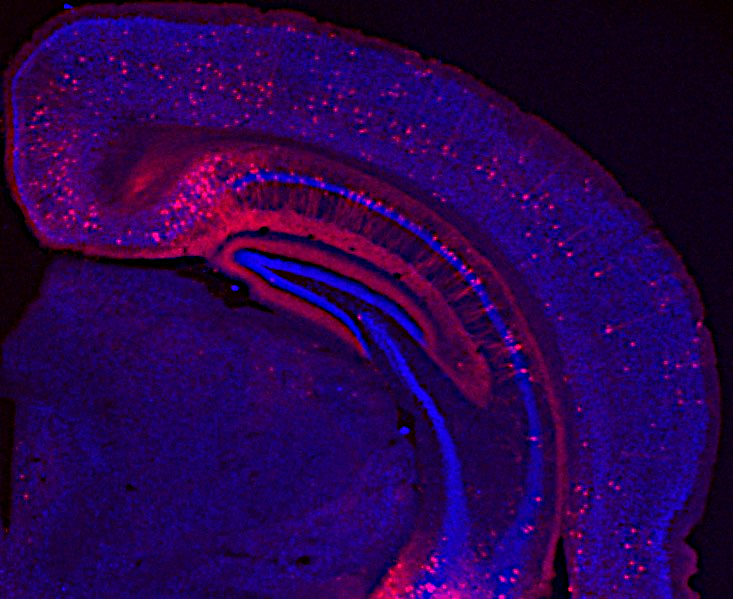

V2

V1

V2

A2

A1

A2

TeA

200μm

**A**

**NeuN**

**mCherry**

**B**

**B**

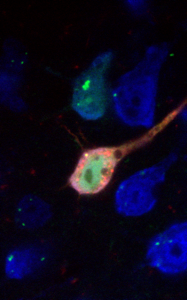

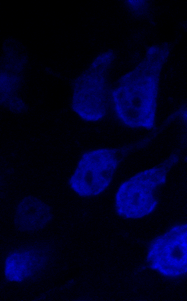

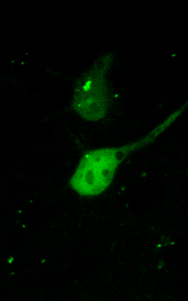

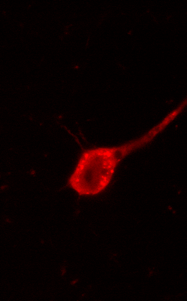

10μm

**C**

**NeuN**

**GFP**

**mCherry**

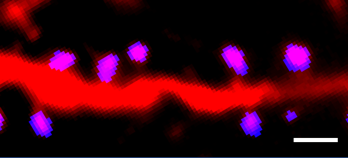

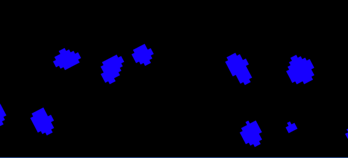

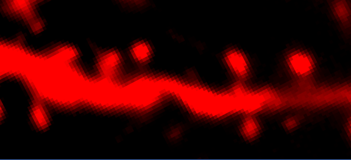

**D**

**mCherry**

**Mask**

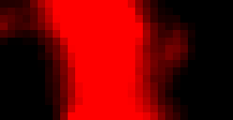

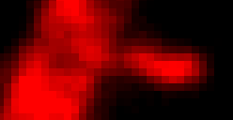

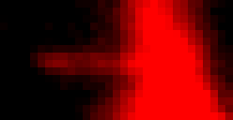

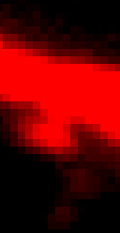

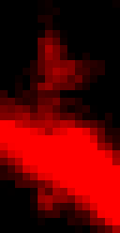

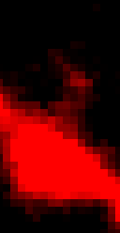

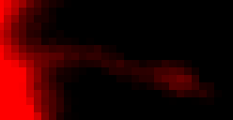

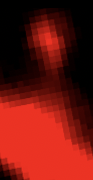

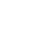

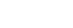

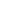

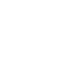

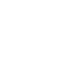

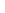

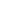

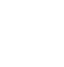

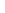

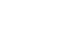

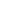

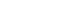

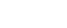

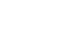

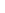

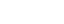

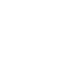

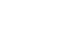

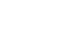

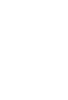

Short stubby

Long stubby

Thin

Short mushroom

Long mushroom

Branched

Atypical

Filopodia

**E**

1μm

1μm

1μm

**Supplemental Figure 2** Neuron sampling and image processing methods. **A)** 1.25x 2-D image of ROI including five regions of sensory cortex of a male CN mouse in the study. NeuN (blue) and mCherry but not GFP fluorescence visible at 1.25x. Dashed lines designate atlas-determined region boundaries and white box in V1 outlines 10x image of L5 pyramidal cells shown in **B**. Scalebar = 200μm. **B)** 10x 2-D image of L5 pyramidal cells with GFP (top) and mCherry (middle) fluorescence. White carets point out the two L5 pyramidal cells with the highest GFP+mCherry+ (merge, bottom) fluorescent intensity. Scalebar = 50μm. **C)** 60x 2-D image of GFP+mCherry+ pyramidal cell from Ms8-164. White box outlines the cell body of the L5 pyramidal cell of interest. Merge far right. Scalebar = 10μm. **D)** Masking strategy used to calculate dendritic spine object volume. For two neurons per mouse, each spine included in neuron DSD was manually marked in Slidebook6. Then, each marked spine object was manually masked using the brush tool in Slidebook6 in the 3D planes in which it occurred. Far left panel shows a 2-D image (single z-plane) of an unmasked representative dendritic segment with dendritic protrusions from Neuron 3 of Ms6-114. Middle panel shows the mask only for the protrusions on this dendritic segment. Far right panel shows the merge. Scalebar = 1μm. **E)** Dendrite protrusion category examples. Scalebar = 1μm.

**Supplemental Figure 3**

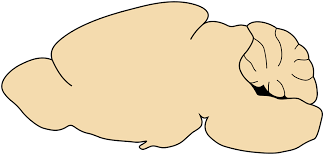

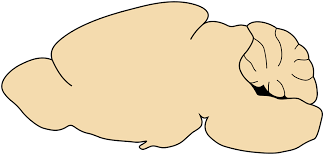

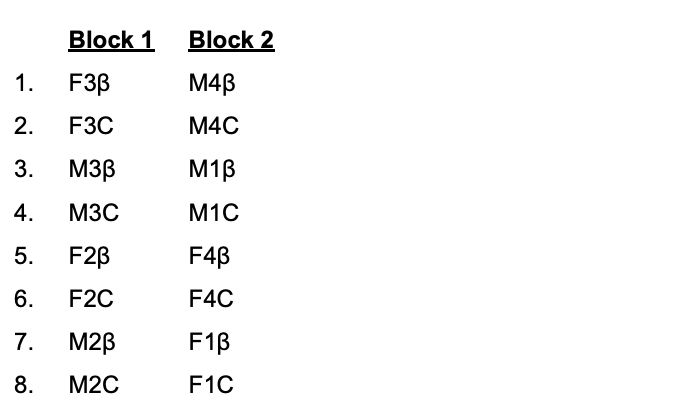

Tissue collection, homogenate preparation, RNAse treatment, BCA

**C**

Randomize order, IP in 2 blocks, add SIL β4 to each sample

**D**

Reorder, S-TRAP PIC creation, trypsin digestions

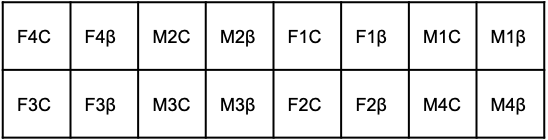

PIC creation, trypsin digestions

**E**

Final reorder, MS

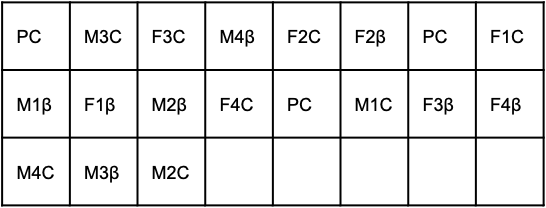

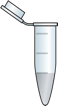

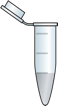

**B**

**A**

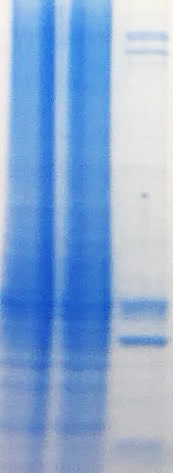

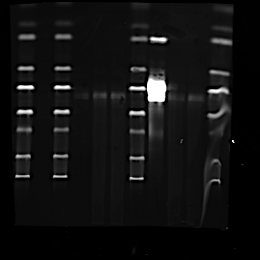

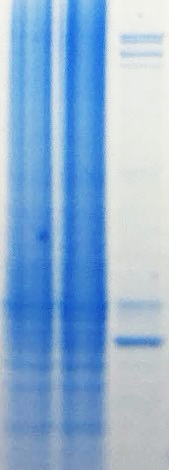

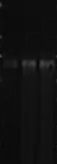

Input

Sup

IP

**β4**

Input

Sup

IP

**CN**

250

50

25

**β4**

250

50

25

**β4**

**Supplemental Figure 3** Co-IP and CoIP-MS methods. **A)** (Above) β4 was immunoprecipitated from RNAse treated mouse brain lysate (n=1 adult female C57Bl/6J mouse) using 10μg mouse anti-Ca_V_β4 antibody (Neuromab #75-054) per 1mg beads (Input=lysate prior to antibody coupling, Sup=supernatant, IP=immunoprecipitant) run out on a gel. Non-antibody coupled beads were used as negative control (CN). β4 isoforms are present at 37.5-59kD (predominant isoforms at ~55kD). (Below) β4 and β4-IP proteins using anti-Ca_V_β4 antibody (Neuromab #75-054) and detection with a goat anti-CACNB4 antibody (Everest Biotech #EB06591, 1:1000) and donkey anti-goat 800 secondary antibody (LiCor #926-32214, 1:10,000). CoIP-MS study design in **B-E**. **B)** For each brain, bilateral cerebral cortex and underlying structures were separated from the olfactory bulb at the optic chiasm, and tissue posterior to the cerebral cortex, midbrain junction. Mouse brain tissue was homogenized using Triton x-100 lysis buffer, treated with RNAse and total protein amount determined using BCA. **C)** Sample name abbreviations were created e.g. Female mouse #3 β4-IP (F3β), Male #1 CN (M3C), and immunoprecipitation (IP) run in two blocks with sample pairs in random order. 1μL stable isotope labeled (SIL) β4 protein was added to each sample (samples=16). Note: sample tubes were number coded starting during this step to blind experimenter to group. Sample name abbreviations rather than number codes are used in schematic to increase clarity. **D)** Samples were reordered, and S-TRAP performed. A pooled instrument control (PIC) was created from all samples and each sample digested with trypsin. **E)** Samples were reordered a final time and loaded onto the LC-MS for analysis.

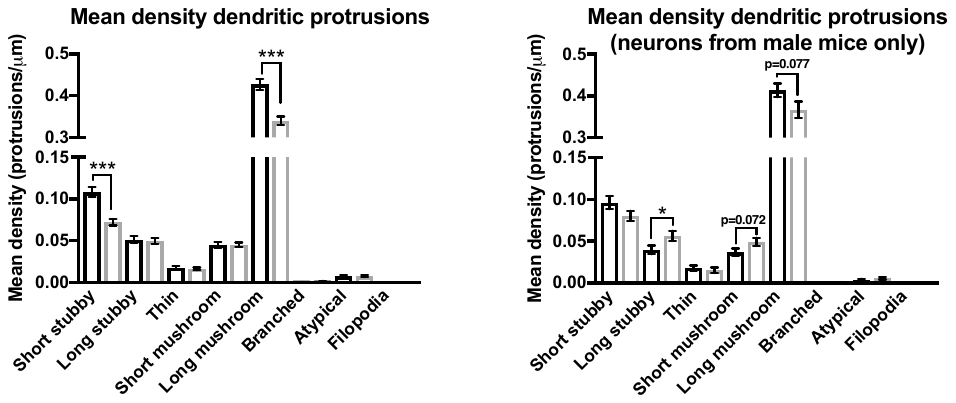

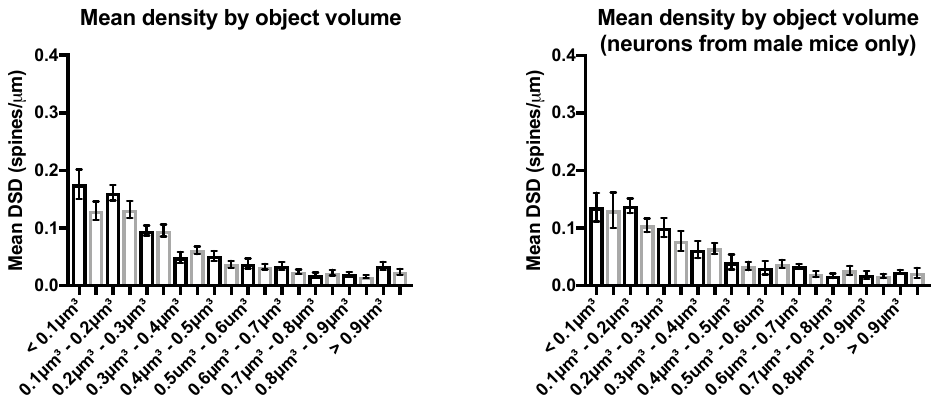
**Supplemental Figure 4**

**B**

**A**

**Supplemental Figure 4** Impact of β4OE on dendritic spines of male mice. **A)** Unlike in females, in male mice β4OE did not significantly reduce mean DSD of masked objects with small < 0.1μm^3^ volume (F=0.017, DF=1, p=0.900). Genotype also did not significantly alter mean DSD of masked objects of 0.1μm^3^ - 0.2μm^3^ volume in male mice (F=3.620, DF=1, p=0.81). Error bars = SEM. **B)** β4OE significantly increased mean density of long stubby spines in male mice (F=4.202, DF=1, p=0.044). There was a trend level increase in mean density of short mushroom (F=3.332, DF=1, p=0.072) and a trend level decrease in mean density of long mushroom (F=3.215, DF=1, p=0.077) spines in male β4OE mice. Error bars = SEM.

**Supplemental Table 1**

| **Peptide** | **Protein ID** | **β4-IP Mean** | **Negative CN Mean** | **Log2(β4-IP)-Log2(CN)** | **p value** |
| --- | --- | --- | --- | --- | --- |
| ILLDAGFTNELVQNYWSK | CA2D1 | 4679428.52 | 430883.648 | 3.44096209 | 0.0020032 |
| HLNVQIAASEK | CACB1 | 1972810.91 | 575075.819 | 1.7784286 | 0.000678 |
| EGGDIAFIPSPQR | CACB3 | 12280054.2 | 1034063.14 | 3.56992075 | 0.00132655 |
| TSLAPIIVHVK | CACB4 | 220800749 | 4296093.28 | 5.68357594 | 6.6278E-06 |
| GSSAGFLTLHNAFPK | CCG8 | 26893944.7 | 2476365.74 | 3.44098507 | 2.2347E-05 |
| GPDGEPQPGLESQGR | CAC1B | 3660770.58 | 984309.532 | 1.89496339 | 0.07737068 |
| SKTDLLNPEEAEDQLADIASVGSPFAR | CAC1A | 7684689.21 | 1429661.12 | 2.42631369 | 0.00037217 |
| SLFIFGEDNIVR | CAC1E | 15626148.1 | 1367039.48 | 3.51483538 | 4.2502E-05 |
| HASLDGASPYFK | CBARP | 7077283.81 | 386774.592 | 4.19363085 | 0.00011575 |
| LADFGLAIEVEGEQQAWFGFAGTPGYLSPEVLR | KCC2A | 16655492.9 | 4292426.25 | 1.9561328 | 0.00029843 |
| NSLVSPAQEPAPLQTAMEPQTTVVHNATDGIK | KCC2G | 11039219.1 | 3554233.61 | 1.6350277 | 0.00211111 |
| WQNVHFHCSGAPVAPLQ | KCC2B | 48924331.4 | 8320871.35 | 2.55574562 | 0.00031296 |
| VTEQLIEAINNGDFEAYTK | KCC2D | 170897637 | 51071238.1 | 1.74254951 | 9.0052E-05 |
| HIFALFNTEQR | DYN1 | 1337059441 | 248763365 | 2.42621766 | 5.726E-05 |
| NLVDSYVAIINK | DYN2 | 26011838.4 | 4424921.46 | 2.55544462 | 0.00036523 |
| DFINSELLAQLYSSEDQNTLMEESAEQAQR | DYN3 | 4692986.01 | 844595.315 | 2.47417401 | 3.1317E-06 |
| GPGPLQER | MATR3 | 9081529.79 | 1012921.79 | 3.16441255 | 0.00030925 |
| ELIFEETAR | MK01 | 19980737 | 5684580.29 | 1.81348406 | 0.00049902 |
| YAGLTFPK | MK10 | 47936716.2 | 11171215.2 | 2.10134495 | 0.0006719 |
| EIQILLR | MK03 | 26591656.5 | 5242495.99 | 2.34264789 | 0.00012184 |
| NIIGLLNVFTPQK | MK08 | 18472179 | 3011726.26 | 2.61669151 | 6.4344E-05 |
| KLIHLEIKPAIR | MP2K1 | 7366876.12 | 699448.954 | 3.3967623 | 0.00061777 |
| LQGTHYSVQSDIWSMGLSLVELAIGR | MP2K2 | 9130170.09 | 595500.023 | 3.93846827 | 4.9442E-05 |
| SDVWSLGITLYELATGR | MP2K4 | 14163070.2 | 1889143.79 | 2.90632961 | 5.4254E-05 |
| ILANGQMNEQDIR | MP2K5 | 1688528.28 | 158518.721 | 3.4130412 | 0.02430607 |
| EAFEQPQTSSTPPR | MP2K6 | 12080013.4 | 1895764.2 | 2.67177063 | 0.00155127 |
| DVKPSNILLDER | MP2K7 | 21391015.4 | 4721868.32 | 2.17957525 | 0.0001709 |
| SLLAAQQTFVDR | PK3C3 | 14133065.9 | 2561537.4 | 2.4639926 | 0.0003328 |
| TLVTGGATPELEALIAEENALR | KCMA1 | 15441800 | 4596895.79 | 1.74810907 | 0.00029363 |
| KGTEELYAIK | KPCA | 17227148.3 | 3350753.57 | 2.36212641 | 3.603E-05 |
| DIKEHAFFR | KPCB | 125821888 | 30482936.7 | 2.04530912 | 0.00024614 |
| SEEEAKFPTMNR | KPCD | 3634260.09 | 746119.948 | 2.28418219 | 0.00059478 |
| LAAGAESPQPASGNSPSEDDRSK | KPCE | 25555749.4 | 3750681.52 | 2.76842323 | 8.7931E-06 |
| LVLASIDQADFQGFTYVNPDFVHPDAR | KPCG | 119358003 | 34747156.5 | 1.78032847 | 8.6122E-05 |
| DVVLMDDDVECTMVEK | KPCT | 33955244.7 | 3786334.84 | 3.16476053 | 0.02319497 |
| YFAQEALTVLSLA | 2AAA | 60900968.3 | 17238564.7 | 1.82082551 | 7.5425E-05 |
| FFEEPEDPSSR | 2ABD | 13957147.7 | 3499596.63 | 1.99574359 | 0.00284384 |
| INLWHLAITDR | 2ABG | 7969074.15 | 1967713.85 | 2.01789169 | 7.9454E-05 |
| EAPVPRPTPQVAASGGQS | 2A5B | 5852125.35 | 1087748.13 | 2.42761614 | 0.00019822 |
| ELFEGAVR | REM2 | 30209737.4 | 13006994.4 | 1.21572601 | 0.00392079 |
| YVDEAHQYILEFDGGSR | RYR2 | 11191113.9 | 2322597.47 | 2.26854259 | 7.9859E-05 |
| EINDLAESGAR | RYR3 | 161878090 | 1147912.95 | 7.13975068 | 8.677E-05 |
| ESALILLQTVPK | ZNT1 | 4402207.23 | 1475242.63 | 1.5772748 | 0.01102076 |
| SLFTEPSEPLPEEPK | ZNT3 | 33314252.1 | 4630270.12 | 2.84697124 | 8.3469E-05 |
| LTELLESDPSVR | ZNT9 | 4013997.57 | 744806.691 | 2.43010181 | 0.06512629 |
| LLTNLGLGEIK | S39AA | 8379085.3 | 1143707.33 | 2.87307485 | 0.00157128 |
| NTLNPYYNESFSFEVPFEQIQK | SYT1 | 126552581 | 15797161.5 | 3.00199967 | 8.6181E-05 |
| VPMNTVDLGQPIEEWR | SYT2 | 16821234.5 | 2471263.78 | 2.76696267 | 0.00022299 |
| YDYESETLIVR | SYT6 | 12489316.5 | 2271135.4 | 2.4592089 | 4.5682E-05 |
| IYLLPDKK | SYT7 | 5776414.96 | 764432.873 | 2.91771266 | 2.751E-05 |
| GELQVSLSYQPVAQR | SYT11 | 5802165.07 | 1491297.49 | 1.96002326 | 0.003395 |
| VSLLPDEQIVGISR | SYT12 | 45324978.2 | 7494428.56 | 2.59641594 | 2.6405E-05 |
| TGSVEAQTALK | SYT13 | 17223089.9 | 3194783.8 | 2.43055378 | 0.00041057 |
| QLLQTDVSQGSDPFVK | SYT17 | 7371700.44 | 2053695.22 | 1.84377535 | 0.00040559 |

**Supplemental Table 1**

Table with VGCCs and other previously reported β-interacting proteins identified within the β4 interactome of male and female P84 mice. In nearly all cases, multiple peptides per Protein ID were identified. Peptide with the lowest significant p value per Protein ID is reported. As expected, the VGCC subunits Ca_V_2.1, Ca_V_2.2, Ca_V_2.3 and α_2_δ1 were identified in the list of protein/peptides detected using the CoIP LC-MS approach utlized in the current study. Furthermore, ryanodine receptors 1 and 2 were identified in the present as well as previously published studies as were dynamin 1-3, phosphoinositide 3-kinase (PI3K), calcium/calmodulin-dependent kinase II (CaMKII, protein ID label KCC2), members of the synaptic protein family synaptotagmin, zinc transporter 1 (plus zinc transporter 3, 9 and 10), members of the Mitogen-activated protein kinase (MAPK) and MAPK kinase families (protein ID labels MK and MP2K), members of the Protein Kinase C family of proteins and the relatively recently discovered and described β-anchoring and-regulatory protein (BARP)^1-13^.

SUPPLEMENTAL REFERENCES

1	Cheng, W., Altafaj, X., Ronjat, M. & Coronado, R. Interaction between the dihydropyridine receptor Ca2+ channel β-subunit and ryanodine receptor type 1 strengthens excitation-contraction coupling. *Proceedings of the National Academy of Sciences* **102**, 19225-19230 (2005).

2	Buraei, Z. & Yang, J. The β subunit of voltage-gated Ca 2+ channels. *Physiological Reviews* **90**, 1461-1506 (2010).

3	Gonzalez-Gutierrez, G., Miranda-Laferte, E., Neely, A. & Hidalgo, P. The Src homology 3 domain of the β-subunit of voltage-gated calcium channels promotes endocytosis via dynamin interaction. *Journal of Biological Chemistry* **282**, 2156-2162 (2007).

4	Sheng, Z.-H., Westenbroek, R. E. & Catterall, W. A. Physical link and functional coupling of presynaptic calcium channels and the synaptic vesicle docking/fusion machinery. *Journal of Bioenergetics and Biomembranes* **30**, 335-345 (1998).

5	Levy, S. *et al.* Molecular basis for zinc transporter 1 action as an endogenous inhibitor of L-type calcium channels. *Journal of Biological Chemistry* **284**, 32434-32443 (2009).

6	Beharier, O. *et al.* Crosstalk between L-type calcium channels and ZnT-1, a new player in rate-dependent cardiac electrical remodeling. *Cell Calcium* **42**, 71-82 (2007).

7	Ohana, E. *et al.* Silencing of ZnT-1 expression enhances heavy metal influx and toxicity. *Journal of Molecular Medicine* **84**, 753-763 (2006).

8	Segal, D. *et al.* A role for ZnT-1 in regulating cellular cation influx. *Biochemical and Biophysical Research Communications* **323**, 1145-1150 (2004).

9	Fitzgerald, E. M. The presence of ca2+ channel β subunit is required for mitogen‐activated protein kinase (mapk)‐dependent modulation of α1b ca2+ channels in cos‐7 cells. *The Journal of Physiology* **543**, 425-437 (2002).

10	Béguin, P. *et al.* BARP suppresses voltage-gated calcium channel activity and Ca2+-evoked exocytosis. *Journal of Cell Biology* **205**, 233-249 (2014).

11	Nakao, A. *et al.* Comprehensive behavioral analysis of voltage-gated calcium channel beta-anchoring and-regulatory protein knockout mice. *Frontiers in Behavioral Neuroscience* **9**, 141, doi:10.3389/fnbeh.2015.00141 (2015).

12	Abiria, S. A. & Colbran, R. J. CaMKII associates with CaV1. 2 L‐type calcium channels via selected β subunits to enhance regulatory phosphorylation. *Journal of Neurochemistry* **112**, 150-161 (2010).

13	Grueter, C. E., Abiria, S. A., Wu, Y., Anderson, M. E. & Colbran, R. J. Differential regulated interactions of calcium/calmodulin-dependent protein kinase II with isoforms of voltage-gated calcium channel β subunits. *Biochemistry* **47**, 1760-1767 (2008).
